# TIRAP drives myelosuppression through an Ifnγ-Hmgb1 axis that disrupts the marrow microenvironment

**DOI:** 10.1101/2020.02.26.967018

**Authors:** Rawa Ibrahim, Aparna Gopal, Megan Fuller, Patricia Umlandt, Linda Chang, Joanna Wegrzyn-Woltosz, Jeffrey Lam, Jenny Li, Melody Lu, Jeremy Parker, Aly Karsan

## Abstract

Activation of inflammatory pathways is associated with bone marrow failure syndromes, but how specific molecules impact on the marrow microenvironment is not well elucidated. We report a novel role for the miR-145 target, Toll/Interleukin-1 receptor domain containing adaptor protein (TIRAP), in driving bone marrow failure. We show that TIRAP is overexpressed in various types of myelodysplastic syndromes (MDS), and suppresses all three major hematopoietic lineages.. Constitutive expression of TIRAP in hematopoietic stem/progenitor cells (HSPC) promotes upregulation of *Ifnγ*, leading to bone marrow failure. Myelopoiesis is suppressed through Ifnγ-Ifnγr-mediated release of the alarmin, Hmgb1, which disrupts the marrow endothelial niche. Deletion of *Ifnγ* or Ifnγr blocks Hmgb1 release and is sufficient to reverse the endothelial defect and prevent myelosuppression. In contrast, megakaryocyte and erythroid production is repressed independently of the Ifnγ receptor. Contrary to current dogma, TIRAP-activated Ifnγ-driven marrow suppression is independent of T cell function or pyroptosis. In the absence of Ifnγ, TIRAP drives myeloproliferation, implicating Ifnγ in suppressing the transformation of bone marrow failure syndromes to myeloid malignancy. These findings reveal novel, non-canonical roles of TIRAP, Hmgb1 and Ifnγ function in the marrow microenvironment,and provide insight into the pathophysiology of preleukemic syndromes.

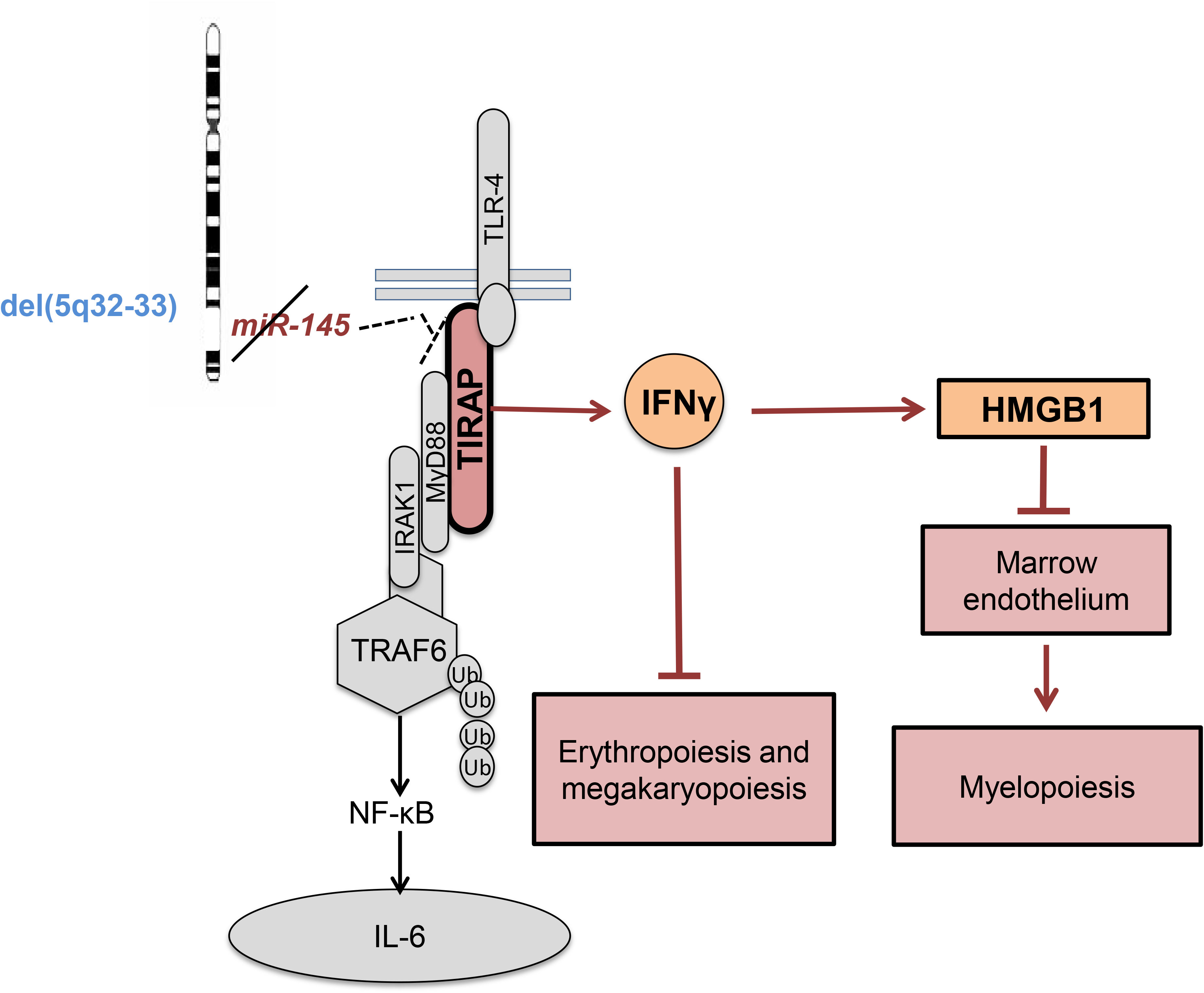
Graphical Abstract: Model of proposed mechanism of TIRAP-induced BMF. Constitutive TIRAP expression in marrow cells releases Ifnγ, which directly impacts on megakaryocyte and erythroid production, but indirectly suppresses myelopoiesis through the release of the alarmin, Hmgb1, which disrupts the marrow endothelial compartment.

## Introduction

Bone marrow failure (BMF) syndromes are a group of clinically and pathologically heterogeneous disorders that result in peripheral blood cytopenias. They are broadly divided into acquired and inherited conditions. Myelodysplastic syndromes (MDS), a type of BMF syndrome, are a group of hematopoietic stem cell malignancies, which are characterized by dysplastic morphology, cellular dysfunction and peripheral blood cytopenias. MDS patients also have a significantly increased risk of transformation to acute myeloid leukemia (AML) (1). The heterogeneity of BMF disorders is evident from the large spectrum of mutations associated with them (2–6).

Dysregulation of immune responses has been implicated in MDS and other BMF disorders (5, 7–10). A pro-inflammatory milieu and sensitivity of hematopoietic stem/progenitor cells (HSPC) to inflammatory cytokines such as TNF-α, IL-6 and IFNγ have been shown to suppress normal hematopoiesis (5, 11). In particular, there is significant evidence for IFNγ playing a key role in marrow failure syndromes (12–14). However, there are different theories, which are sometimes conflicting, regarding the mechanism by which IFNγ promotes BMF. On the one hand IFNγ has been suggested to recruit T cells that mediate cytopenias through immune destruction of hematopoietic cells. On the other hand, T cells have been suggested to produce IFNγ which is postulated to result in death of marrow cells (15, 16). More recently, IFNγ has been shown to inhibit thrombopoietin (TPO) and possibly erythropoietin (EPO) signaling through their cognate receptors by forming heterodimeric complexes and interfering with ligand-receptor interactions, thereby leading to impaired megakaryocytic and erythroid differentiation (17,18).

However, this does not explain the myelosuppression that is also seen with excess IFNγ (19). Recent evidence suggests a role for alarmin-triggered pyroptosis, a Caspase-1-dependent proinflammatory lytic cell death, in mediating myelosuppression in MDS (20). Alarmins such as S100A8 and S100A9 have been suggested to drive pyroptotic cell death in MDS (20–24). S100A8 and S100A9 have also been shown to cause an erythroid differentiation defect in Rps14-haploinsufficient HSPC in a cell-autonomous manner (22). However, the generality of alarmin-triggered BMF across MDS is not known. Given the heterogeneity of MDS, other alarmins may also be involved, but have not been studied thus far, and whether IFNγ can also trigger alarmin-induced death is not known.

There is also very little information on the triggers that underlie upregulation of IFNγ in these marrow failure syndromes (12–14, 23). Interstitial deletion of chromosome 5q is the most common cytogenetic abnormality observed in MDS, accounting for approximately 10% of all cases (25). Our lab has previously shown that miR-145, which is located on chromosome 5q, targets TIRAP, but the role of TIRAP in marrow failure has not been studied (5). In this study, we identify a novel role for the innate immune adaptor protein TIRAP in dysregulating normal hematopoiesis through induction of *Ifnγ*. TIRAP induces *Ifnγ* through a non-canonical pathway, and leads to the development of BMF. TIRAP-induced activation of Ifnγ releases the alarmin, Hmgb1, which suppresses the marrow endothelial niche, which in turn promotes myeloid suppression. In contrast, erythroid and megakaryocytic suppression appears to be a direct effect of Ifnγ that is independent of the Ifnγ receptor. In contrast to what has previously been suggested for the mechanism of Ifnγ-induced marrow failure, BMF induced by the TIRAP-Ifnγ-Hmgb1 axis does not require NK or T cell function nor pyroptosis.

## Results

### Constitutive expression of TIRAP results in BMF

To examine TIRAP expression across MDS subtypes, we analyzed the results of a gene expression study performed on CD34^+^ cells from MDS patients and controls (26). Although TIRAP expression was increased in del(5q) MDS as expected, a subset of patients diploid at chromosome 5q also showed increased expression suggesting a broader role for TIRAP in MDS (Fig. 1a). To determine whether constitutive expression of TIRAP contributes to BMF, we stably expressed TIRAP or an empty vector control in mouse HSPC (Fig. 1b). Lethally irradiated recipient mice were transplanted with TIRAP-transduced HSPC along with wild-type helper cells. Marrow engraftment of mice transplanted with TIRAP-expressing HSPC was comparable to controls (Fig. 1c). However, peripheral blood output of TIRAP-transplanted mice was significantly reduced compared to control mice (Fig. 1c). Mice transplanted with TIRAP-expressing HSPC had significantly reduced overall survival due to BMF compared to control mice (*P* < 0.0001) (Fig. 1d-g). We observed significant anemia (*P* < 0.0001), leukopenia (*P* = 0.007), thrombocytopenia (*P* = 0.003) and splenomegaly in TIRAP-transplanted mice at 4 weeks (Fig. 1e and f). Histological examination of marrow sections and quantification of marrow cellularity revealed significant reduction in total cell number in TIRAP-transplanted marrows compared to controls (*P* < 0.001) (Fig. 1g; Supplemental Fig. 1a).

**Fig. 1:**
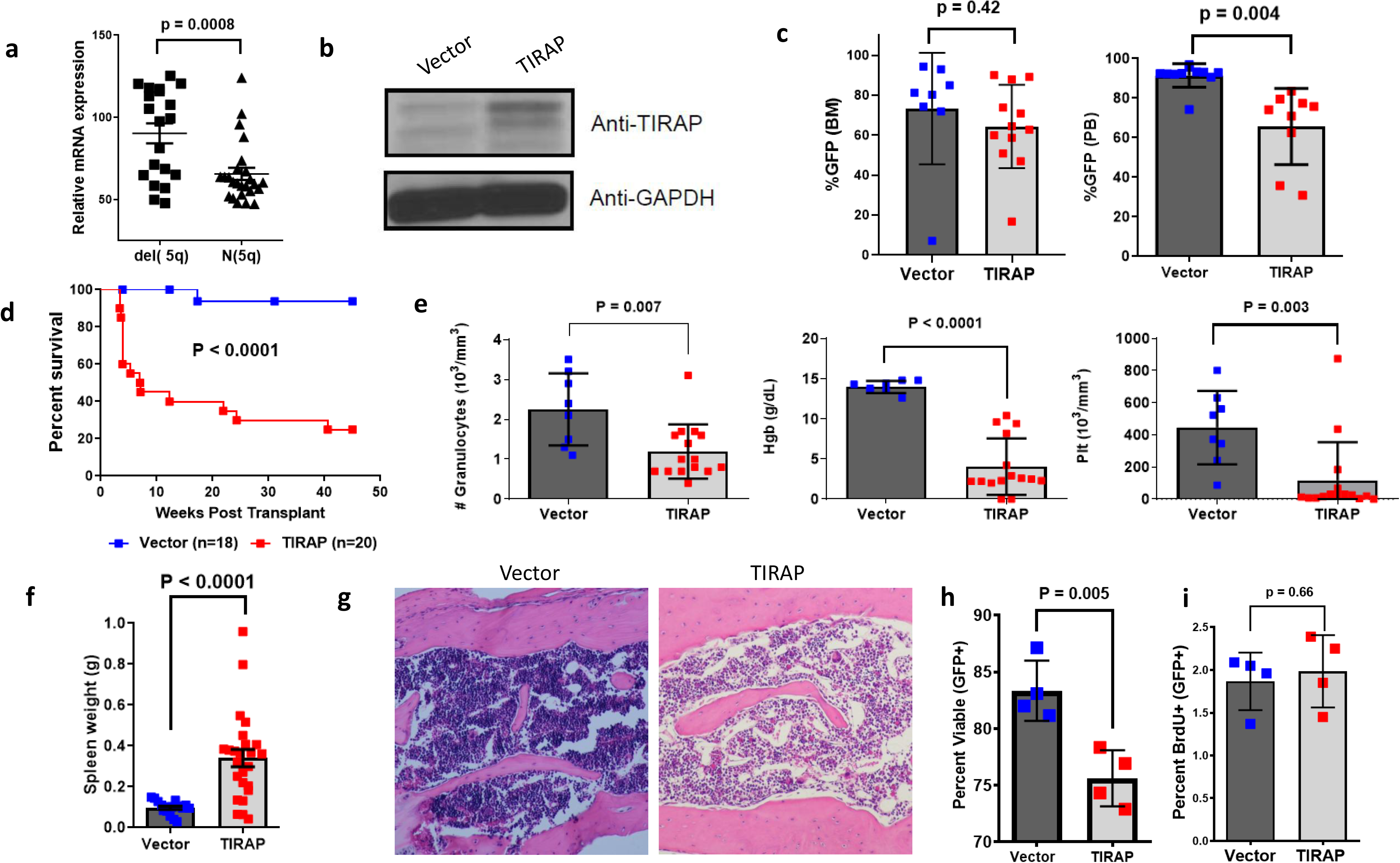
Constitutive expression of TIRAP in hematopoietic stem/progenitor cells leads to BMF. (a) TIRAP mRNA expression in CD34^+^ cells isolated from del(5q) patient marrow and marrow with normal diploid copy number (N) at chromosome 5q (from data generated in ^22^). (b) Wildtype mouse HSPC transduced with empty vector (Vector) or TIRAP were sorted for GFP expression and immunoblotted with antibodies against TIRAP and GAPDH. (c) Bone marrow and peripheral blood engraftment in wild-type mice transplanted with Vector- or TIRAP-transduced marrow cells. (d) Kaplan-Meier survival curves for wild-type mice reconstituted with wild-type HSPC transduced with Vector (n = 18) or TIRAP (n = 20). (e) Granulocyte counts, hemoglobin (Hgb) and platelet counts from TIRAP-transduced (n = 15) and Vector-transduced wild-type mice (n = 8) at experimental end-point. (f) Spleen weights of Vector and TIRAP-transplanted wild-type mice at experimental end-point. (g) H&E-stained marrow sections showing marrow hypocellularity in TIRAP transplanted wild-type mice compared to Vector. (h) Percent viable (Annexin V^-^/PI^-^) HSPC from Vector (n = 4) or TIRAP (n = 4) transplanted wildtype mice at 3-4 weeks post-transplant. (i) BrdU incorporation was measured in wild-type HSPC transduced with TIRAP (n = 4) or Vector (n = 4).

Bone marrow homing experiments revealed no difference in the homing ability nor the clonogenic activity of TIRAP-transduced HSPC compared to controls (Supplemental Fig. 1b-d), suggesting that the BMF was not due to a homing defect. There was increased cell death but no difference in proliferation *in vivo* in TIRAP-expressing HSPC compared to control (Fig. 1h).

### Constitutive TIRAP expression has cell non-autonomous effects on hematopoiesis

Examination of the marrow of moribund mice showed significant impairment of TIRAP-transplanted mice to reconstitute hematopoiesis (Supplemental Fig. 2). This impairment was evident despite transplantation of sufficient wild-type helper cells to sustain normal hematopoiesis, suggesting that BMF subsequent to TIRAP expression was likely due to cell non-autonomous mechanisms. Differentiation into mature granulocytes (Gr-1^+^) was compromised in TIRAP-transduced HSPC compared to that of controls (Supplemental Fig. 2b). Interestingly the myeloid defect was more pronounced in the non-transduced population and extended to the early granulocyte/macrophage population as evidenced by a 50% reduction in the CD11b^+^Gr-1^+^ population (Supplemental Fig. 2c). Erythroid differentiation of transduced HSPC as well as wildtype competitor cells was also significantly impaired in TIRAP-transplanted mice, with the impairment more evident in the co-transplanted wild-type cells (Supplemental Fig. 2d-f), while megakaryocytic reconstitution was only impaired in wild-type cells (Supplemental Fig. 2g-i). These findings suggest that cell non-autonomous mechanisms play a large role in TIRAP-mediated BMF, and that there is relative protection within the TIRAP-expressing population.

### TIRAP alters the marrow microenvironment

In contrast to our findings *in vivo* (Fig. 1h and i) TIRAP-expressing HSPC cultured *in vitro* displayed increased viability and proliferation compared to control HSPC suggesting a requirement of the microenvironment for TIRAP induced BMF (Supplemental Fig. 2j and k). Given the role of immune dysregulation in various BMF syndromes, we examined whether the lymphoid populations in the marrow of the transplanted mice were affected. The analysis revealed a significant reduction in CD19^+^B cells in the marrows of TIRAP-transplanted mice compared to controls (Fig. 2a-c), as has been previously reported in low-risk MDS patients (27). Interestingly, although there was an increase in T cells in TIRAP-expressing marrows compared to control marrows, this was entirely due to increased expansion or recruitment of wild-type (GFP^-^) T cells (Fig. 2a-c).

**Fig. 2:**
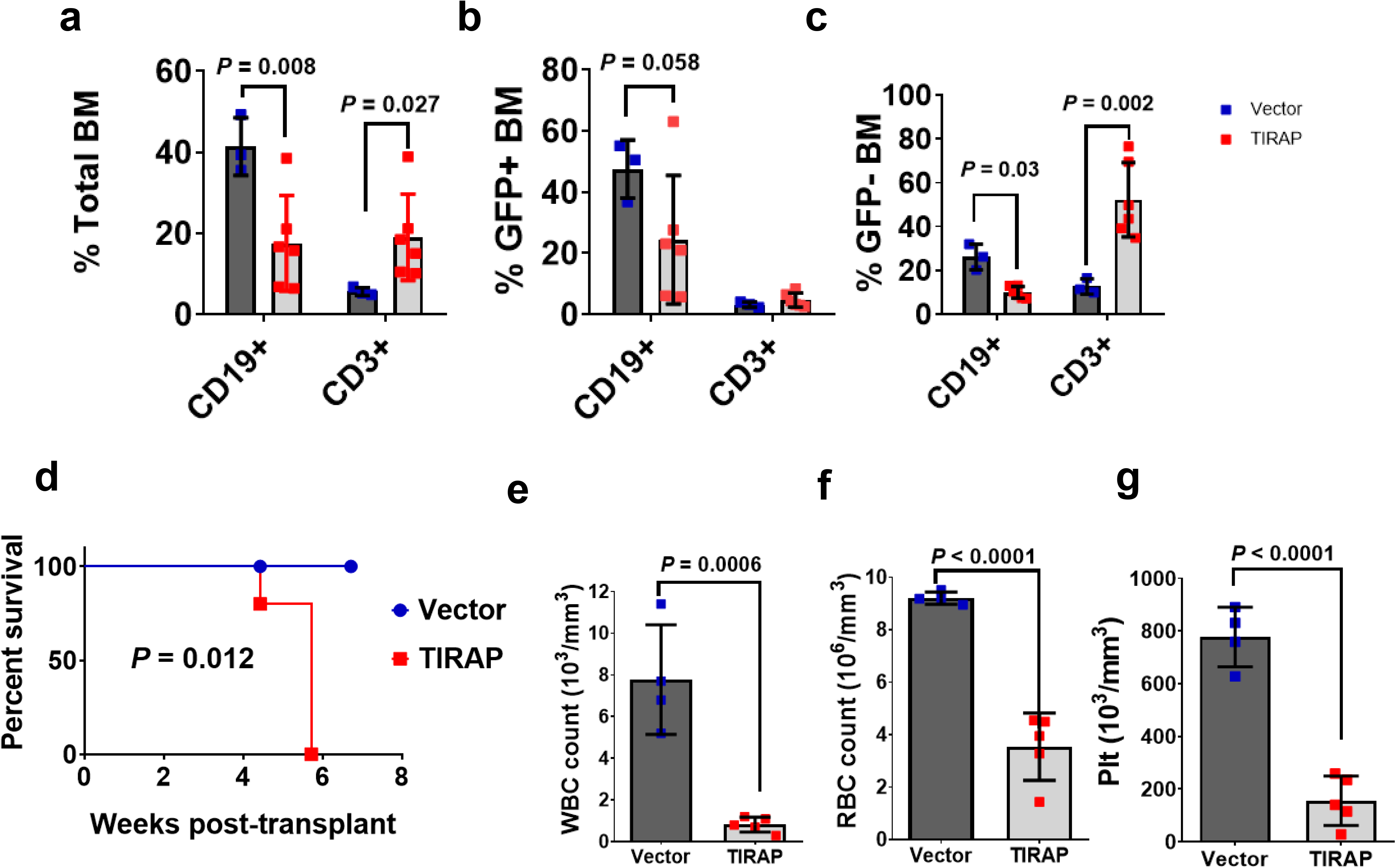
Absence of autologous T cells and NK cells does not prevent marrow failure. Immunophenotyping of lymphoid (a-c) lineages from bone marrow of wild-type mice transplanted with Vector-(n = 3) or TIRAP-(n = 6) transduced wild-type HSPC at 3-4 weeks post transplant. Total HSPC, as well as the contribution of transduced (GFP+) and competitor cells (GFP-) are represented. (d) Kaplan-Meier survival curve for NSG mice transplanted with wild-type HSPC transduced with Vector (n = 4) or TIRAP (n = 5). (e) White blood cell (WBC) counts, (f) red blood cell (RBC) counts, and (g) platelet (Plt) counts in TIRAP (n = 5) and Vector (n = 4) transduced mice at endpoint.

Although T cells have been suggested to be key mediators of marrow cytopenias, these studies have mostly been associative rather than demonstrating direct causation (15, 16, 28). To address whether the expansion or recruitment of effector T cells was required for BMF, we transplanted NSG (NOD/*scid*/gamma) mice, which are deficient in functional T, B and NK cells, with wild-type HSPC transduced with TIRAP or control in the absence of wild-type competing cells to avoid wild-type T cell expansion. Similar to wild-type transplants, NSG mice transplanted with TIRAP-expressing HSPC developed BMF with pancytopenia in a similar timeframe as wild-type mice (Fig. 2d-g) suggesting that T and NK cells are not necessary for the development of BMF.

To determine whether TIRAP-expressing cells altered the ability of marrow stroma to support hematopoiesis, we tested the ability of GFP- or YFP-labeled HSPC to competitively repopulate wild-type mice following exposure to TIRAP or empty vector-conditioned environments (Fig. 3a, Supplemental Fig. 3). Following a 3-week conditioning period, marrows were myeloablated with busulfan, followed by transplantation of GFP- or YFP-labeled wild-type HSPC. After a two week engraftment period, marrow collected from TIRAP and control-conditioned mice were pooled, and one mouse equivalent of the mixed marrow was transplanted into lethally-irradiated wild-type recipients. The contribution of GFP^+^ and YFP^+^ cells to hematopoietic reconstitution in the marrow was determined after 11 weeks (Fig. 3b). The contribution of myeloid cells derived from TIRAP-conditioned marrows was significantly impaired compared to that of cells derived from control-conditioned marrows (4.1% compared to 95.9% *P* < 0.0001). This is consistent with the reduced marrow cellularity observed in mice transplanted with HSPC constitutively expressing TIRAP, suggesting that exposure to TIRAP-expressing cells reduces the capability of the marrow microenvironment to support normal hematopoiesis.

**Fig. 3:**
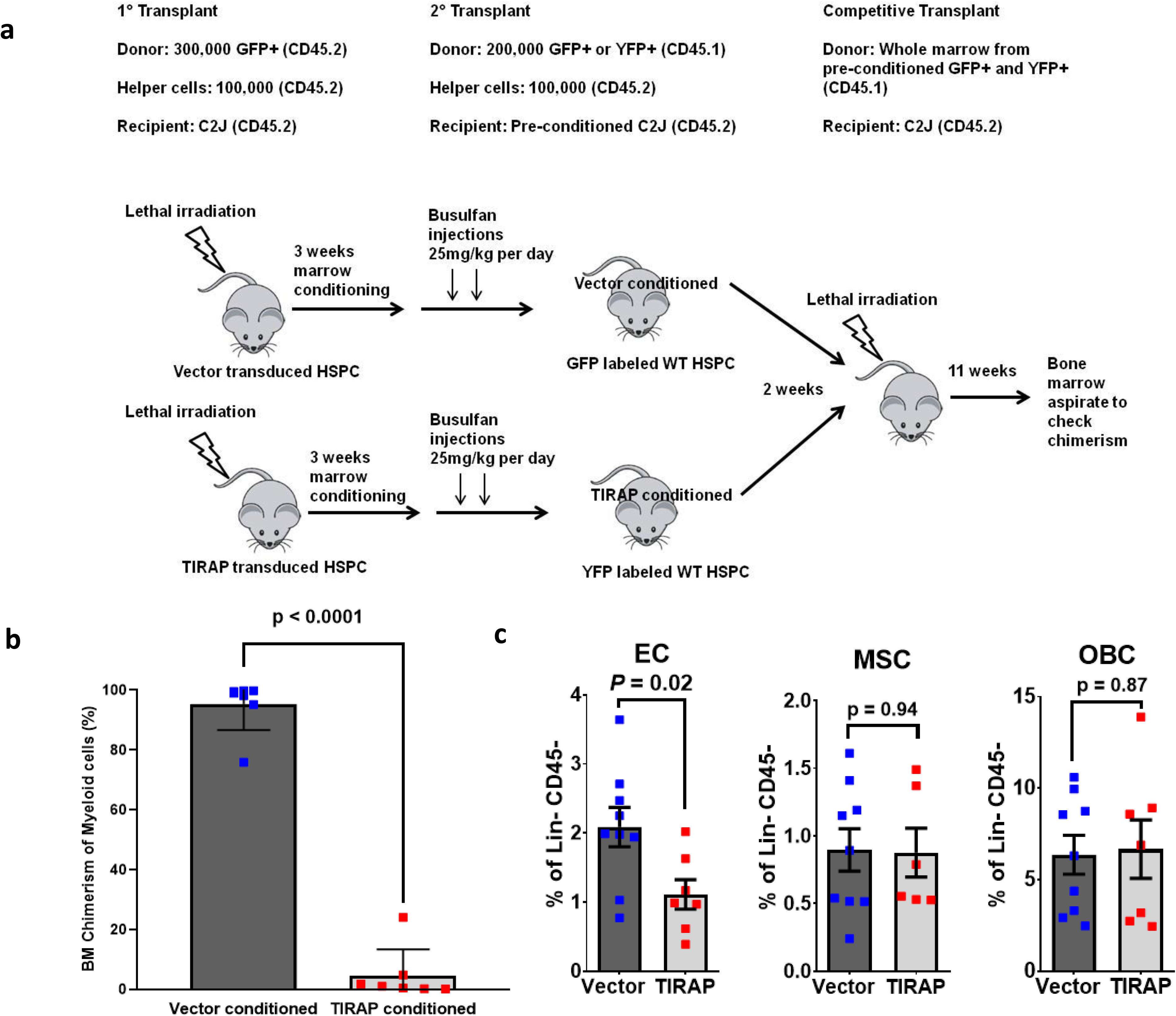
Constitutive TIRAP expression disrupts the marrow microenvironment. (a) Outline of experimental schema. Wild-type mice were transplanted with TIRAP- or Vector-transduced wild-type HSPC (CD45.2) and allowed to condition the marrow microenvironment for 3 weeks. After 3 weeks mice were myeloablated with busulfan. GFP- or YFP-labeled wildtype HSPC (CD45.1) were then transplanted into the myeloablated mice and allowed to engraft for 2 weeks, following which marrow was harvested (Vector-conditioned n = 7; TIRAP-conditioned n = 7), pooled and transplanted competitively into new wild-type transplant recipients (n = 7). (b) GFP and YFP myeloid chimerism in marrow from competitive transplants was measured after 11 weeks (n = 7). (c) Frequency of Lin^-^CD45^-^CD31^+^ endothelial cells, Lin^-^CD45^-^Sca-1^+^CD51^+^ mesenchymal stromal cells, and Lin^-^CD45^-^Sca-1^-^CD51^+^ osteoblastic cells from primary transplanted wild-type mice reconstituted with TIRAP- (n = 7) or Vector-transduced (n = 9) wild-type HSPC.

HSPC are dependent on marrow stromal cells to provide signals for proliferation, differentiation, and survival (24, 29) and several studies have implicated stromal abnormalities in the pathogenesis of myeloid malignancies (24, 30, 31). We thus analyzed the composition of the marrow microenvironment to determine whether TIRAP expression in HSPC alters components of the marrow microenvironment (32) (Supplemental Fig. 4). We found no difference in the percentage of lin^-^CD45^-^Sca-1^+^CD51^+^mesenchymal stromal cells, or in the lin^-^CD45^-^Sca-1^-^ CD51^+^ osteoblast cells derived from TIRAP-transplanted mice compared to controls (Fig. 3c). However, lin^-^CD45^-^ CD31^+^ endothelial cells were found to be reduced by 45% in TIRAP-transplanted mice (Fig. 3c).

### Ifnγ drives TIRAP-induced BMF but limits progression to myeloid malignancy

We postulated that the inhibitory effects of constitutive TIRAP expression on endothelial cells may be due to paracrine factors from TIRAP-expressing cells. To identify soluble factors potentially responsible for the effects on endothelial cells and the subsequent BMF following constitutive TIRAP expression we performed RNA-seq to compare the transcriptional profiles of TIRAP-expressing and vector-transduced HSPC. Using Ingenuity Pathway Analysis we identified possible upstream regulators and found several proinflammatory cytokines to be significantly overrepresented (Supplemental Table 1). Of these, the Ifnγ pathway was identified as the most highly activated pathway in TIRAP-compared to control-transduced HSPC (Supplemental Table 1). Furthermore, Geneset enrichment analysis (GSEA) identified the Ifnγ response as the single most significantly enriched pathway of the Hallmark genesets (Fig. 4a). While most of the enriched gene sets represented proinflammatory pathways, the anti-inflammatory cytokine Il10 was also predicted to be an upstream regulator (Supplemental Table 1). GSEA also revealed enrichment in Il10 signaling in TIRAP-transduced HSPC compared to controls (Fig. 4b).

**Fig. 4:**
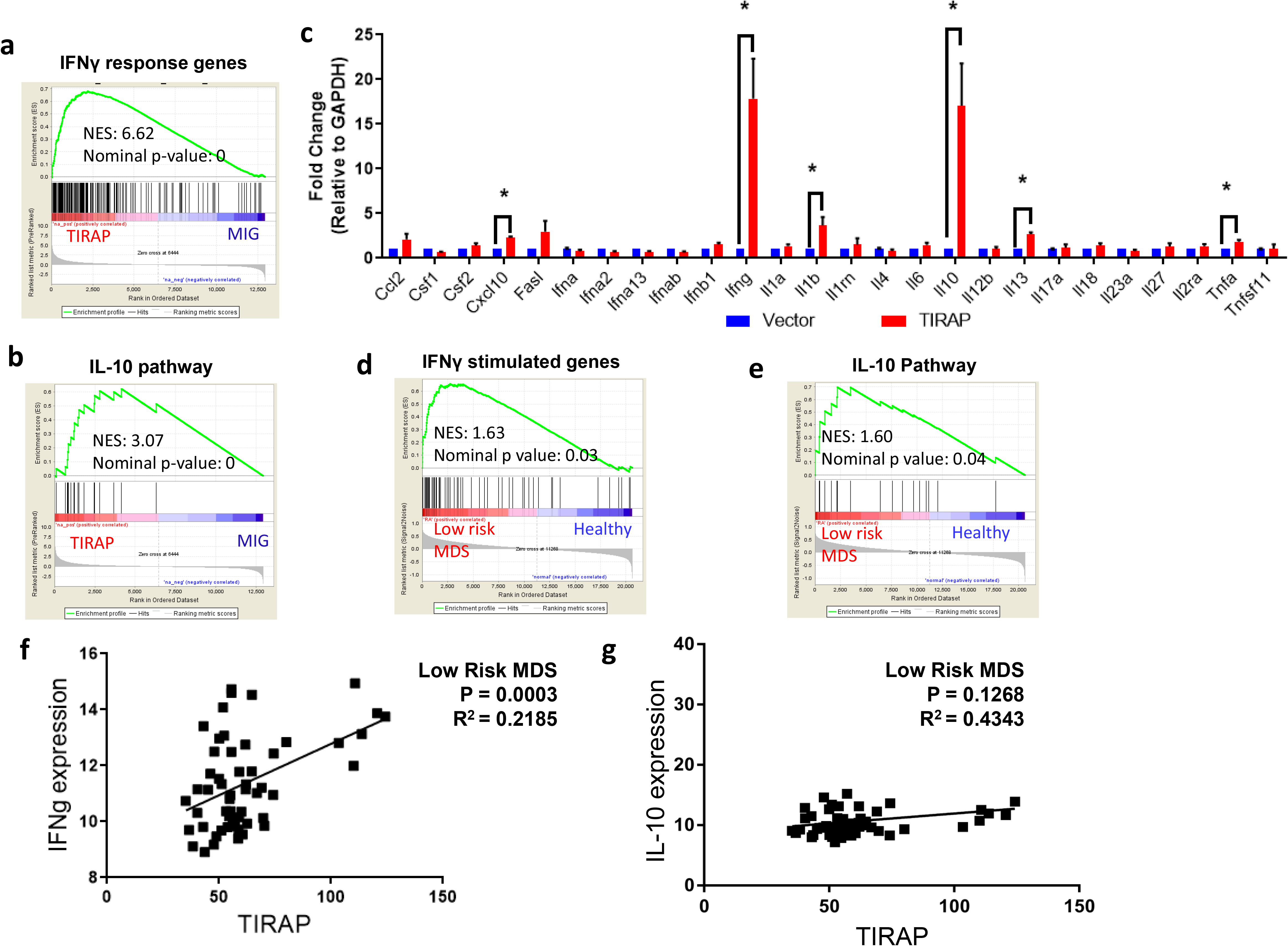
Ifnγ signaling is upregulated following constitutive TIRAP expression in murine HSPC and is activated in low-risk MDS. Geneset enrichment plots for (a) Ifnγ response and (b) Il10 pathway obtained from RNA-seq analysis of wild-type HSPC transduced with TIRAP or Vector. (c) Validation of cytokine gene expression in Vector- and TIRAP-transduced wild-type HSPC by quantitative RT-qPCR. * *P* < 0.05. Geneset enrichment plots from low-risk MDS marrow compared to normal marrow generated for (d) IFNγ stimulated genes, and (e) IL-10 pathway genes. (f) Correlation between IFNγ gene expression and TIRAP gene expression in low-risk MDS patients. (g) Correlation between IL-10 gene expression and TIRAP gene expression in low-risk MDS patients. Data were analyzed from previously published microarray dataset (19) for plots in d-g.

We verified expression of the predicted upstream regulators of differential gene expression and examined other cytokines previously implicated in the pathogenesis of MDS, by quantitative RT-qPCR. *Ifnγ, Il10, Il1b, Il13* and *Tnfa* transcript levels were all increased in TIRAP-expressing HSPC (P < 0.05) (Fig. 4c). As *Ifnγ* and *Il10* showed the greatest differential expression, and were among the top predicted regulators of global changes in gene expression following constitutive TIRAP expression (Fig. 4c), we confirmed that constitutive TIRAP expression resulted in increased concentration of Ifnγ and Il10 in serum of mice transplanted with TIRAP-expressing HSPC and conditioned medium, respectively (Supplemental Fig. 5a,b). Of note, examination of a previously published gene expression dataset (26) showed that *IFNγ* and *IL10* signatures were upregulated in low-risk MDS patients compared to healthy controls (Fig. 4d,e) suggesting clinical significance of these signaling pathways. *TIRAP* expression was also positively correlated with *IFNγ* expression in low-risk MDS patients (*P* = 0.0003; Fig. 4f), but not with *IL10* expression (*P* = 0.1268; Fig. 4g).

To test the functional role of Ifnγ and Il10 in TIRAP-mediated BMF, wild-type recipient mice were transplanted with TIRAP- or control-transduced HSPC from *Ifnγ*^−/−^ or *Il10*^−/−^ donor mice, along with wild-type marrow cells. Mice that received TIRAP-transduced *Ifnγ*^−/−^ HSPC were rescued from BMF, as evidenced by normalized blood cell counts and improved median survival compared to controls (*P* = 0.02; Fig. 5a and b). In contrast, transplantation of TIRAP-expressing *I110*^−/−^ HSPC did not lead to improvement in blood counts nor in overall survival (*P* = 0.9; Fig. 5c and d). Median survival for TIRAP-transplanted mice reconstituted with *Ifnγ*^−/−^ and *IllO*^−/−^ HSPC was 48.6 weeks and 27.2 weeks respectively.

**Fig. 5:**
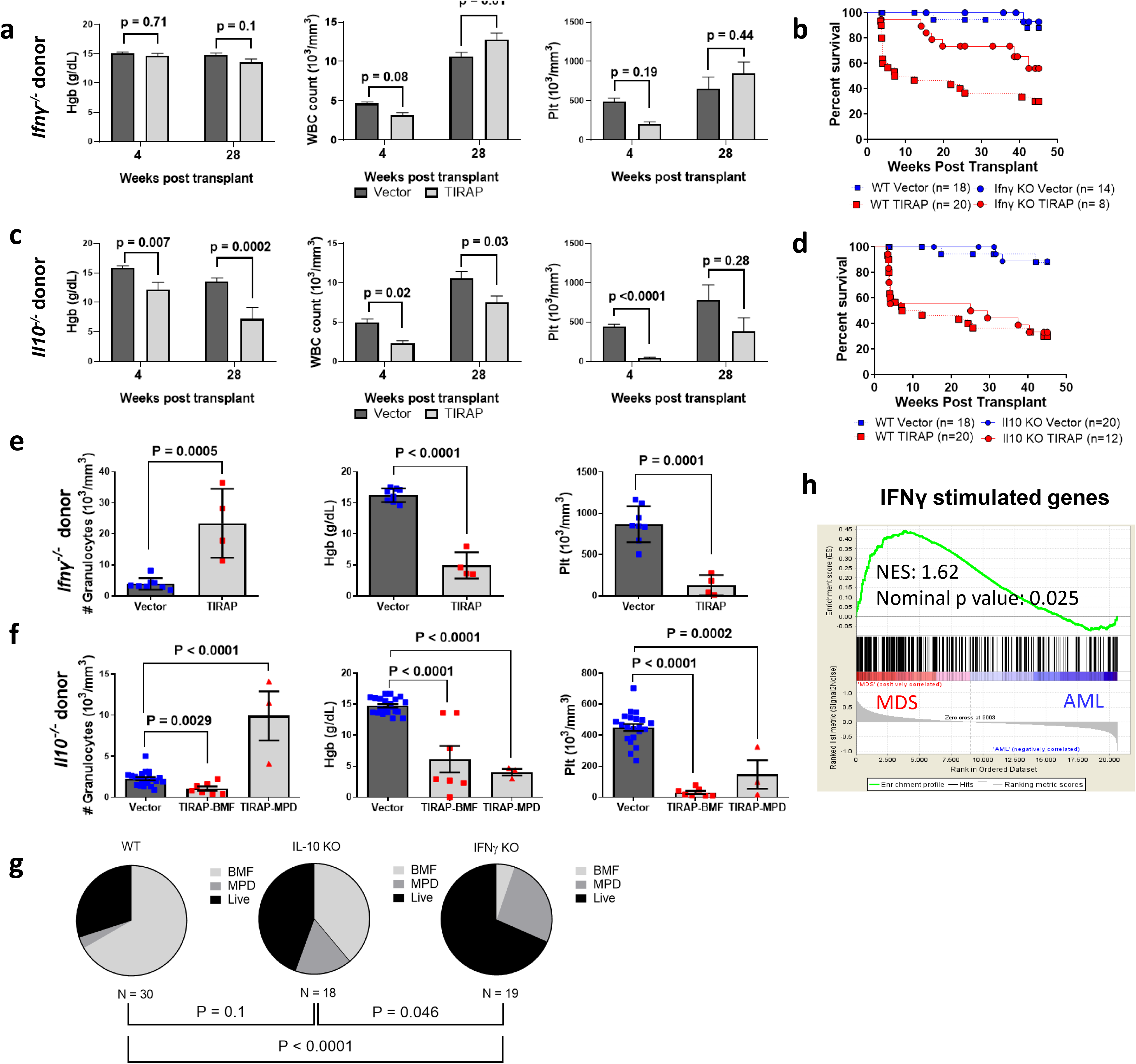
TIRAP requires Ifnγ to promote BMF and inhibit progression to myeloid malignancy. (a) Complete blood counts from mice transplanted with *Ifnγ*^−/−^ HSPC transduced with TIRAP or Vector. (b) Kaplan-Meier survival curves for transplanted wild-type mice reconstituted with HSPC from *Ifnγ*^−/−^ mice transduced with Vector (n = 14) or TIRAP (n = 8), WT vs. *Ifnγ^−/−^, P* = 0.0218. (c) Complete blood counts from wild-type mice transplanted with *Il10*^−/−^ HSPC transduced with TIRAP or Vector. (d) Kaplan-Meier survival curves for primary transplanted mice reconstituted with HSPC from *Il10*^−/−^ mice transduced with Vector (n = 20) or TIRAP (n = 12), WT vs. *Il10*^−/−^, *P* = 0.9197. Survival curves for wild-type mice in panels (b) and (d) are those shown in Fig. 1d. (e, f) Hemoglobin levels, granulocyte counts, and platelet counts at experimental end-point in wild-type mice transplanted with *Ifnγ*^−/−^ or *Il10*^−/−^ HSPC transduced with Vector or TIRAP. (g) Distribution of the difference in cause of mortality between wild-type mice transplanted with WT, *Il10*^−/−^ and *Ifnγ*^−/−^ TIRAP-transduced marrows. Statistical significance was assessed using a Chi square test. (h) GSEA analysis showing enrichment of the IFNγ signature in MDS patients compared to AML patients. BMF: Bone marrow failure, MPD: Myeloproliferative disorder.

As *Ifng* is thought to be produced mainly by T and NK cells (33), the lack of requirement for T and NK cells in driving BMF (Fig. 2) was a surprise. We thus examined which cells express *Ifng* following TIRAP expression. This analysis showed that myeloid and NK cells were the cells most responsible for the increased production of *Ifng* (Supplemental Fig. 5c), thus explaining how the absence of NK cell-derived *Ifng* can be overcome by myeloid-derived *Ifng* to drive TIRAP-mediated BMF in NSG mice.

Although survival of mice transplanted with TIRAP-expressing *Ifnγ*^−/−^ HSPC was extended, these mice still began to die approximately 15 weeks post-transplant (Fig. 5b). Moribund mice presented with leukocytosis, anemia and thrombocytopenia consistent with a myeloproliferative disorder (MPD) (Fig. 5e). No improvement in overall survival was observed for recipients of *I110*^−/−^ HSPC (Fig. 5c and f). In contrast to mice that received TIRAP-transduced *Ifny^-^* HSPC, there was no significant increase in MPD in mice that received TIRAP-transduced *I110*^−/−^ HSPC (Fig. 5g). The significant increase in progression to MPD in the context of Ifnγ deficiency suggests that Ifnγ plays a role in the suppression of both normal and malignant hematopoiesis, and implies that leukemic progression may require suppression of Ifnγ signaling. Indeed, GSEA analysis of gene expression data comparing MDS and AML patients (34) showed enrichment of the IFNγ signature in MDS patients compared to AML (Fig. 5h), suggesting that IFNγ signaling is upregulated in MDS, and that repression of this pathway may be important for leukemic transformation.

### Ifnγ has an indirect effect on marrow stromal cells

As interferons have been shown to be associated with inflammasome formation and pyroptosis (35) and S100 alarmin-driven BMF has been associated with pyroptosis (20–24), we asked whether pyroptosis was activated in this model, by immunoblotting transduced HSPC for Caspase-1. Increased levels of Caspase-1 were noted in TIRAP-transduced HSPC compared to controls, and flow cytometry revealed an increase in activated Caspase-1 in TIRAP-expressing marrow cells from TIRAP-transplanted mice compared to controls (Supplemental Fig. 6a, b). Caspase-1 activation was mainly observed within the myeloid compartment, and was present in both the transduced population as well as the wild-type helper cell population (Supplemental Fig. 6b). However, deletion of Caspase-1 did not attenuate Ifnγ or Il10 expression. (Supplemental Fig. 6c). To determine whether the increase in Caspase-1 activation was required for myelosuppression, we constitutively expressed TIRAP or empty vector in *Casp1*^−/−^ HSPC and transplanted cells into lethally irradiated *Casp1*^−/−^ recipient mice. Loss of Caspase-1 did not reverse TIRAP-induced pancytopenia and mice succumbed to BMF within 4 weeks of transplantation similar to TIRAP-transduced and transplanted wild-type animals (Supplemental Fig. 6d, e and Fig. 1d), suggesting that in this model pyroptosis may be an epiphenomenon rather than the cause of BMF.

To determine whether BMF mediated by TIRAP via Ifnγ was directed through the Ifnγ receptor (Ifnγr), we performed transplants in which *Ifnγr*^−/−^ recipient mice were transplanted with *Ifnγr*^−/−^ HSPC constitutively expressing TIRAP or control, as well as *Ifnγr*^−/−^ helper cells. Unexpectedly, transplantation of TIRAP-expressing *Ifnγr*^−/−^ HSPC resulted in rapid death (Fig. 6a). In contrast to *Ifnγ*^−/−^ transplants (Fig. 5a), erythrocyte and platelet counts did not improve in mice transplanted with TIRAP-expressing *Ifnγr*^−/−^ HSPC, although leukocyte counts were maintained (Fig. 6b). The difference between the *Ifnγ*^−/−^ model and the *Ifnγr*^−/−^ model may be reconciled based on the continued expression of Ifnγ in the *Ifnγr*^−/−^ mice, which may directly interfere with TPO signaling (17,18). As noted, similar to the *Ifnγ^−/−^* model, deletion of Ifnγr resulted in maintenance of the myeloid population (Fig. 6b). All *Ifnγr*^−/−^ mice expressing TIRAP displayed massive splenomegaly (mean spleen weight of 239 mg in TIRAP vs. 89 mg in control mice, *P* = 0.0014) due to myeloid infiltration of the spleen (Supplemental Fig. 7). This finding is reminiscent of the myeloproliferation seen in TIRAP transplants using *Ifnγ*^−/−^ HSPC above. The absence of Ifnγ-sensitive cells in both donor and recipient mice led to maintenance of the marrow endothelial population (Fig. 6c), suggesting that the reduction seen in the endothelial population in wild-type mice transplanted with TIRAP-expressing HSPC is due to Ifnγ and that endothelial cells support the myeloid population, as has been suggested previously (36).

**Fig. 6:**
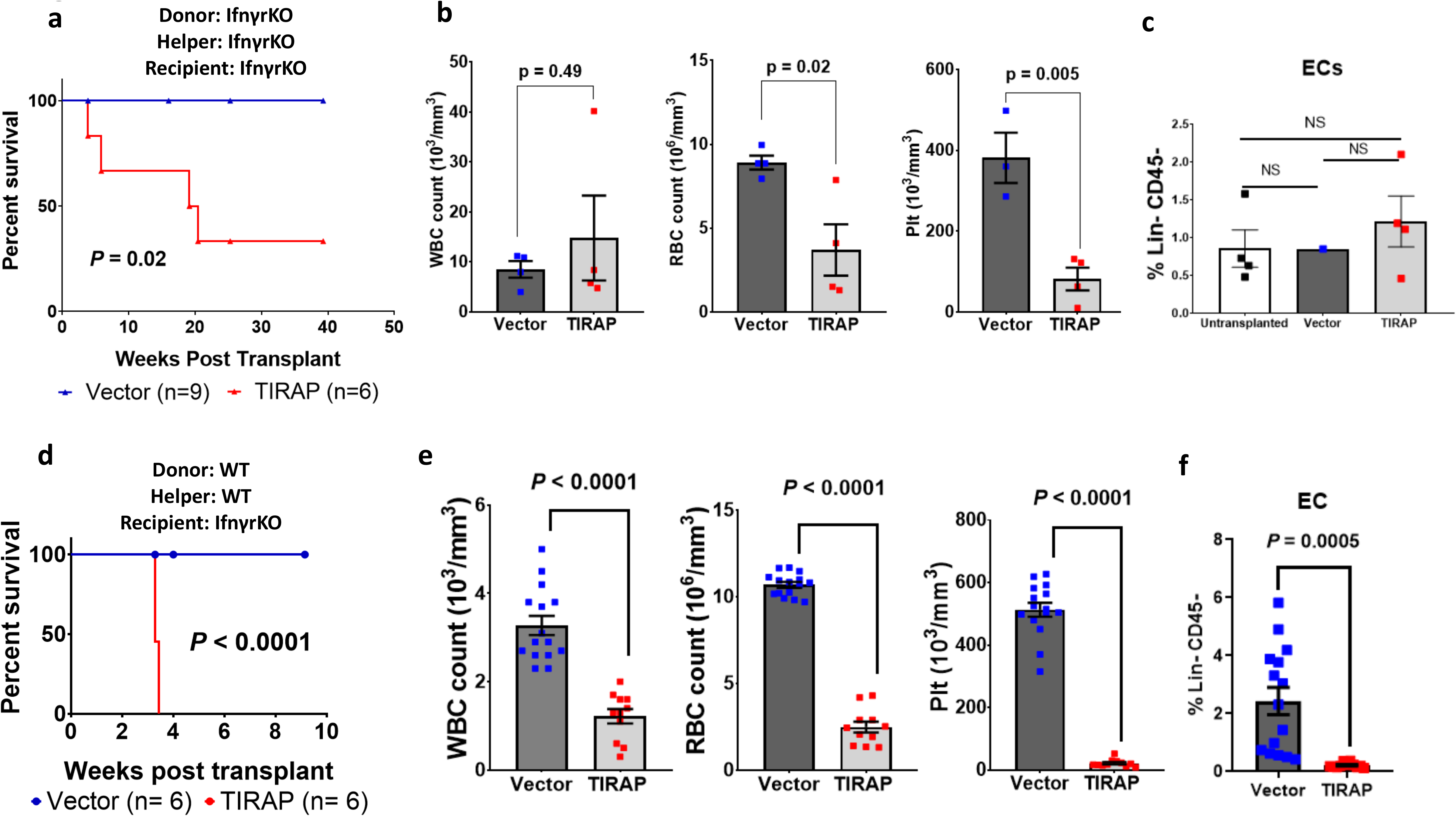
Ifnγ has an indirect effect on marrow endothelial cells. (a) Kaplan-Meier survival curves for *Ifnγr*^−/−^ primary transplanted mice reconstituted with *Ifnγr^−/−^* HSPC transduced with vector (n = 9) or TIRAP (n = 6). (b) Complete blood counts at experimental end-point of *Ifnγr*^−/−^ mice transplanted with *Ifnγr*^−/−^ HSPC transduced with Vector or TIRAP. (c) Frequency of Lin^-^CD45^-^CD31^+^ endothelial cells, Lin^-^CD45^-^Sca-1^+^CD51^+^ mesenchymal stromal cells, and Lin^-^CD45^-^Sca-1^-^CD51^+^ osteoblastic cells from *Ifnγ^−/−^* mice transplanted with TIRAP- or Vector-transduced HSPC. (d) Kaplan-Meier survival curves for *Ifnγ^−/−^* recipient mice reconstituted with wild-type HSPC transduced with Vector (n = 6) or TIRAP (n = 6). (e) Complete blood counts of *Ifnγ^−/−^* mice transplanted with TIRAP- or Vector-transduced wild-type HSPC. (f) Frequency of Lin^-^CD45^-^CD31^+^ endothelial cells in *Ifnγr^−/−^* recipient mice transplanted with TIRAP- or Vector-transduced WT HSPC.

To test whether Ifnγ was directly responsible for the inhibitory effects on marrow endothelial cells, we transplanted *Ifnγr*^−/−^ recipient mice with wild-type HSPC transduced with either TIRAP or control as well as wild-type helper cells. Surprisingly, these mice had no improvement in overall survival and developed BMF characterized by anemia, thrombocytopenia, and leukopenia, similar to wild-type recipients, despite the insensitivity of marrow stromal cells to Ifnγ (Fig. 6d,e). Furthermore, the marrow endothelial population was depleted, suggesting an indirect effect of Ifnγ on the marrow endothelium (Fig. 6f).

### Hmgb1 acts downstream of Ifnγ to deplete marrow endothelium and suppress myelopoiesis independent of pyroptosis

We examined the RNA-seq data comparing TIRAP-expressing HSPC to vector-transduced HSPC (Supplemental Table 2), to identify signaling pathways downstream of Ifnγ potentially responsible for the endothelial defect. Although pyroptosis is not a cause of BMF in our model (Supplemental Fig. 6), in line with recent work studying the role of alarmins in mouse and human BMF syndromes (20–24), we found the Hmgb 1 pathway to be significantly activated in TIRAP-expressing HSPC (Fig. 7a,b, Supplemental Table 3). We confirmed that Hmgb1 was present in the supernatant of TIRAP-expressing HSPC one day post-transduction (Fig. 7c). To determine whether Hmgb1 was released downstream of Ifnγ, we transduced *Ifnγ*^−/−^ or wild-type HSPC with TIRAP, and assayed for Hmgb1 in the supernatant. As shown, in Fig. 7d, Hmgb1 release was blocked in *Ifnγ*^−/−^ HSPC expressing TIRAP. On the other hand, Hmgb1 release was not blocked in *Casp1*^−/−^ HSPC (Fig. 7d), suggesting pyroptosis-independent release of Hmgb1 in this model of TIRAP-mediated BMF.

**Fig. 7:**
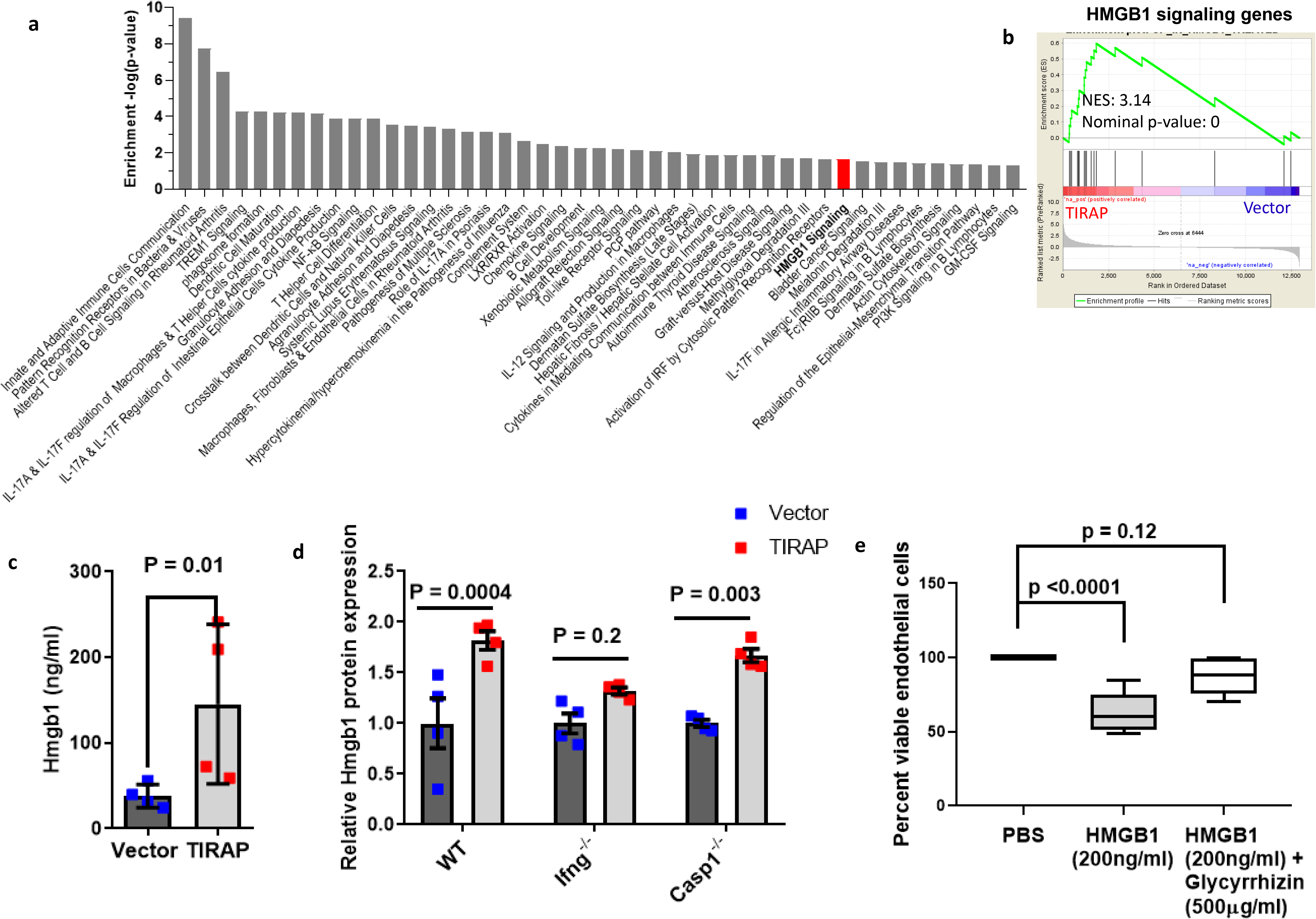
Hmgb1 acts downstream of TIRAP and Ifnγ to deplete marrow endothelium and suppress myelopoiesis. (a) Ingenuity pathway analysis showing canonical pathways predicted to be upregulated following constitutive TIRAP expression in mouse HSPC. The dashed line indicates the significant cut off value of 1.3, which represents log(p = 0.05). (b) GSEA showing enrichment in Hmgb1 signaling in HSPC expressing TIRAP compared to Vector. (c) ELISA for Hmgb1 in supernatant collected from wild-type HSPC expressing TIRAP or Vector. (d) Relative induction of Hmgb1 in wild-type, *Ifnγ*^−/−^ and *Casp1^−/−^* HSPC transduced with TIRAP or Vector, as determined by ELISA of supernatant of Vector- or TIRAP-transduced cells. (e) Percent viability of HUVEC after 24 h treatment with Hmgb1 with or without Glycyrrhizin.

To study the role of Hmgb1 on endothelial cell depletion, we treated human endothelial cells, with either Hmgb1 alone or Hmgb1 and its inhibitor, Glycyrrhizin. There was a significant reduction in the viability of the Hmgb1-treated endothelial cells by 24 hrs compared to PBS (control) treated cells (Fig. 7e). Treatment with Glycyrrhizin restored the viability of the endothelial cells (Fig. 7e) implicating a cytotoxic effect of Hmgb1 on endothelial cells.

## Discussion

Dysregulation of the immune system has been reported in all subtypes of MDS (37). Aberrant activation of various components of the innate immune pathway, such as TLRs and IRAK1, has been implicated in altering normal hematopoiesis (7,8,37). In this study we show that aberrant expression of the innate immune adaptor protein TIRAP dysregulates normal hematopoiesis. The constitutive expression of TIRAP in HSPC has cell non-autonomous effects on other HSPC and the microenvironment leading to the development of BMF, characterized by myeloid, megakaryocytic and erythroid suppression. Based on our results we propose a model of BMF in which aberrant TIRAP overexpression in hematopoietic cells releases Ifnγ, which indirectly suppresses myelopoiesis through the release of the alarmin, Hmgb1, but directly impacts on megakaryocyte and erythroid production. Hmgb1 disrupts the marrow endothelial compartment which suppresses myelopoiesis. Thus, the TIRAP-Ifnγ-Hmgb1-mediated BMF is due to two different mechanisms, and contrary to current dogma, both mechanisms are independent of T cell function or pyroptosis.

Previous studies have focussed on immune and inflammatory activation of canonical TLR4-TRAF6-NFκB activation in MDS (5, 7, 8). Activation of TRAF6, an E3 ubiquitin ligase, has been shown to account for neutropenia and thrombocytosis as well as progression to AML (5). In del(5q) MDS, the upregulation of TRAF6 is due to haploinsufficiency of miR-146a, located on the long arm of chromosome 5. However, miR-146a is only lost in about half the patients with del(5q) MDS (38), as it is not located in the minimally deleted region. In contrast, miR-145 is present within the commonly deleted region of chromosome 5q universally deleted in del(5q) MDS (5). TIRAP, a target of miR-145, lies upstream of TRAF6 and is required for TLR2 and TLR4-dependent activation of TRAF6 (5). Interestingly our data show that TIRAP activates *Ifnγ* but not *Il6*, while TRAF6 expression induces *Il6* but not *Ifnγ*, suggesting that BMF induced by constitutive TIRAP expression is independent of canonical TRAF6-NFκB mediated signaling (5). Further, while TRAF6 overexpression leads to leukemia in half the cases (5), constitutive TIRAP expression does not cause myeloproliferation unless Ifnγ is blocked.

Dysregulation of the marrow microenvironment due to genetic aberrations in stromal cells has been shown to lead to ineffective communication between Hematopoietic stem cells (HSC) and the surrounding microenvironment, contributing to the inability of the marrow to sustain normal hematopoiesis (24, 39). In line with these studies, we demonstrate the role of the hematopoietic microenvironment specifically in maintaining normal myelopoiesis. Interestingly, in our model we do not see any effect on mesenchymal stromal cells and osteoblasts. Rather, we see a reduction in the endothelial cell component of the microenvironment. HSC have been shown to reside in close proximity to the marrow sinusoidal endothelium, and endothelial cells are essential for the maintenance of HSC and promoting normal hematopoiesis (40, 41). Consistent with previous data our results demonstrate the crucial role that endothelial cells play in supporting the myeloid population within the marrow (36).

IFNγ has been implicated in the pathophysiology of BMF syndromes (12, 13). A previous study has shown that IFNγ augments the expression of Fas and other pro-apoptotic genes and causes destruction of hematopoietic cells via Fas-mediated apoptosis (14). This study suggested that the destruction of the functionally active HSC, upon exposure to IFNγ, is immune mediated through activated cytotoxic T cells (14). Additionally, it has been suggested that immune-mediated destruction of HSC leads to BMF in aplastic anemia (9). However, in the current study in the context of immune-deficient NSG mice, the lack of functional T, B and NK cells did not attenuate TIRAP-mediated BMF. In our model BMF, although mediated by IFNγ, is not solely due to T cell-mediated immune destruction of the HSPC, suggesting a context-dependent role of IFNγ in regulating hematopoiesis. We cannot rule out that T cells are involved in BMF, but our data suggest that they are not necessary, as Ifnγ derived from myeloid cells is sufficient to drive the phenotype.

Adding to the pleiotropic role of IFNγ, the negative impact of IFNγ on HSPC survival and proliferation has also been attributed to the direct interaction of IFNγ with TPO and possibly EPO (17, 18). IFNγ was shown to bind to TPO and perturb TPO-c-MPL interactions; thereby inhibiting TPO induced signaling pathways (17). TPO is known to promote megakaryocytic and erythroid differentiation (42). Thus the interaction of IFNγ with TPO may explain the lack of rescue in RBC and platelet counts in TIRAP-transplanted mice that lack the Ifnγ receptor.

While Ifnγ appears to directly affect the erythroid and megakaryocyte populations, we noted an indirect effect on the endothelial population of the marrow microenvironment. This indirect effect occurs via Hmgb1, which we show to be elaborated downstream of Ifnγ. Hmgb1 is a non-histone chromatin binding protein and a member of the alarmin family (43, 44). It is known to function as a mediator of inflammatory processes by binding to its receptors including RAGE and a subset of TLRs (44). HMGB1 has been shown to be present in MDS marrow plasma at greater than three times the levels seen in normal marrow plasma (45, 46), similar to the levels seen in the TIRAP model presented here, providing functional relevance to the role of HMGB1 in BMF syndromes.

Alarmins, in particular S100A8 and S100A9, have been shown to direct NLRP3 inflammasome-mediated activation of Caspase-1, which has been suggested to lead to pyroptotic cell death of HSPC in MDS (20). This mechanism has been proposed to be cell non-autonomous where S100A9 in “non-malignant” myeloid-derived suppressor cells activates inflammasome-mediated pyroptosis of HSPC in MDS (20, 21). However, S100A8 and S100A9 have also been shown to cause an erythroid differentiation defect in Rps14-haploinsufficient HSPC in a cell-autonomous manner (22). Although HMGB1 has been shown to be present at high levels in MDS marrow plasma, the role of this alarmin in the disease phenotype has not been studied (45, 46). Our data with TIRAP-induced BMF suggests that Caspase-1 mediated pyroptosis is not a driving factor for the observed BMF, as TIRAP expression in *Casp1*^−/−^ cells did not rescue TIRAP-induced BMF or mortality. Rather our findings suggest that Ifnγ-stimulated Hmgb1 release occurs in different cell types in a cell-autonomous and non-autonomous manner. Thus, although Hmgb1 is involved in mediating BMF, activation of Caspase-1 is likely an epiphenomenon that is not causative of the BMF seen in this model.

The current study highlights a non-canonical effect of aberrant TIRAP expression on the endothelial cell component of the marrow microenvironment. We further show a cell non-autonomous and non-pyroptotic role of the alarmin Hmgb1 on the marrow endothelial compartment to mediate the observed myelosuppression. As Hmgb1 is a targetable molecule (44,45), further understanding of the mechanisms by which it affects endothelial cells and myelopoiesis would open up avenues for developing new therapies for BMF.

## Methods

### Mice

Pep3b (Ly5.1), C57Bl/6J-TyrC2J (Ly5.2), and NSG (NOD *scid* gamma - NOD.*Cg-Prkdc^scid^ Il2rg^tm1Wjl^*/SzJ) mice were originally obtained from The Jackson Laboratory and were bred and maintained at the Animal Resource Centre of the British Columbia Cancer Research Centre. Ifnγ^−/−^ (B6.129S7-Ifng^tm1Ts^/J) (stock # 002287), Il-10^−/−^ (B6.129P2-Il10^tm1Cgn^/J) (stock# 002251), *Casp1*^−/−^ (B6N.129S2-Casp1tm1Flv/J) (stock #016621), and Ifnγr^−/−^ mice (B6.129S7-Ifngr1tm1Agt/J) (stock #003288) mice were purchased from The Jackson Laboratory. All strains were maintained in-house at the BC Cancer Research Centre Animal Resource Centre.

### Retroviral vectors, packaging cell lines

Flag-TIRAP was PCR amplified from pCMV2-flag-TIRAP (a gift from Ruslan Medzhitov’ lab) to include *XhoI* and *EcoRI* restriction sites. The amplified Flag-TIRAP was cloned into the MSCV-IRES-GFP (MIG) retroviral vector after digesting with *XhoI* and *EcoRI.* Virus packaging and infection of ecotropic packaging cell lines (GP^+^E86) was performed as previously described^5^ to obtain stable MIG-TIRAP lines. Sequencing information confirmed the expression of human TIRAP isoform b in the MIG-TIRAP constructs and cell lines.

### Bone marrow transplants

8-12 weeks old donor mice were injected i.v. with 5-fluorouracil (150mg/kg) and bone marrow was harvested after four days. Marrow cells were retrovirally transduced with MIG or MIG-TIRAP, and GFP positive cells were sorted. Lethally irradiated (810 rads) recipient mice were transplanted i.v. with 300,000 GFP positive cells and 100,000 non-transduced helper cells. Irradiated mice were given ciprofloxacin/HCl in their drinking water for one month following transplantation. Peripheral blood counts were determined at 4-week intervals using the scil Vet abc Blood Analyzer. When mice became moribund as defined by the Animal Resource centre rules, they were sacrificed and processed to obtain the peripheral blood via cardiac puncture, marrow and spleen at endpoint.

### Niche conditioning transplants

Bone marrow transplants were performed as described above. After three weeks of marrow conditioning, 50 mg/kg of Busulfan was administered i.p. over two days and mice were allowed to recover from weight loss for two days following the last injection. 200,000 GFP or YFP labeled marrow cells along with 100,000 helper cells were then injected i.v. into mice with TIRAP or MIG pre-conditioned marrows. GFP and YFP expressing cells were allowed to engraft for 2 weeks. Marrow cells were then collected from the TIRAP and MIG conditioned mice, mixed together, and one mouse-equivalent of marrow was then transplanted into lethally irradiated recipients. GFP and YFP chimerism in the peripheral blood was then monitored.

### Flow cytometry

For immunophenotypic analysis, we washed and resuspended bone marrow cells in PBS containing 2% calf serum, followed by primary monoclonal antibody (phycoerythrin (PE)- or allophycocyanin (APC)-labelled) staining overnight, followed by analysis on a BD FACSCalibur flow cytometer. Antibodies used were PE-conjugated anti-mouse Gr1 (clone RB6-8C5; BD), APC-conjugated anti-mouse Mac1 (clone M1/70; BD), PE-conjugated anti-mouse CD3 (clone 17A2; BD), APC-conjugated anti-mouse CD19 (clone 1D3; BD), PE-conjugated anti-mouse CD45R/B220 (clone RA3-6B2; BD), PE-conjugated anti-mouse CD4 (clone GK1.5, BD), APC-conjugated anti-mouse CD8a (clone 53-6.7; BD), PE-conjugated anti-mouse CD41 (clone MWReg30; BD), PE-conjugated anti-mouse CD71 (clone C2; BD), and APC-conjugated anti-mouse Ter119 (clone Ter119; Biolegend). Antibodies used for progenitor staining were FITC-conjugated anti-mouse CD45.1 (clone A20; eBioscience), PerCP-Cy5.5-conjugated anti-mouse Gr1 (clone RB6-85C; BD) PerCP-Cy5.5-conjugated anti-mouse Ter119 (clone Ter119; eBioscience), PerCP-Cy5.5-conjugated anti-mouse B220 (clone RA3-6B2; eBioscience), PerCP-Cy5.5-congugated anti-mouse CD3 (clone 172A; BD), PerCP-Cy5.5-conjugated anti-mouse CD4 (clone RM4-5; BD), PerCP-Cy5.5-conjugated anti-mouse CD8a (clone 53-6.7; BD), PerCP-Cy5.5-conjugated anti-mouse IL-7R (clone A7R34; eBioscience), APC-conjugated anti-mouse c-Kit (clone 2B8; eBioscience), APC-Cy7-conjugated anti-mouse CD16/32 (clone 2.42; BD), PE-Cy7-conjugated anti-mouse Sca1 (clone 7; BD), biotinylated anti-mouse CD34 (cloneRAM34; eBioscience), and Streptavidin-PE-TxRed (BD Pharmigen). Cells were analyzed on BD FACS Aria III. Antibodies used for stromal staining were PerCP-Cy5.5-conjugated anti-mouse Ter119 (clone Ter119; eBioscience), PerCP-Cy5.5-conjugated anti-mouse B220 (clone RA3-6B2; eBioscience), PerCP-Cy5.5-congugated anti-mouse CD3 (clone 172A; BD), PerCP-Cy5.5-conjugated anti-mouse CD4 (clone RM4-5; BD), PerCP-Cy5.5-conjugated anti-mouse CD8a (clone 53-6.7; BD), PerCP-Cy5.5-conjugated anti-mouse IL-7R (clone A7R34; eBioscience) PerCP-Cy5.5 conjugated anti-mouse CD5 (Clone 53-7.3; eBioscience), PerCP-Cy5.5 conjugated anti-mouse CD11b (clone M1/70; BD Pharmigen), PE-conjugated anti-mouse CD31 (clone MCC 13.3; BD Pharmingen), APC-Cy7 conjugated antimouse CD45 (clone 30-F11; Biolegend), PE-Cy7-conjugated anti-mouse Sca1 (clone 7; BD), biotinlyated anti-mouse CD51 (clone RMV-7; BD Pharmingen), and Streptavidin-APC (eBioscience).

### Apoptosis assay

For Annexin V staining, 2 x 10^5^ cells were washed in PBS, and resuspended in 100ul Annexin V binding buffer (10 mM HEPES; 140 mM NaCl; 2.5 mM CaCl_2_; pH 7.4) and APC-conjugated Annexin V (1:50) and propidium iodide (1:1000) was added. Following 15 min incubation, an additional 200ul Annexin V binding buffer was added, and samples were analyzed by flow cytometry.

### BrdU proliferation assay

Transduced 5FU-treated cells were serum starved overnight in DMEM + 2% FBS followed by treatment with BrdU (1 μM) for 4 hours at 37°C and 5% CO_2_. Transplanted marrow cells were serum starved overnight in DMEM + 2% FBS, followed by a media change to DMEM + 10% FBS and treatment with BrdU (1μM) for 4 hours at 37°C and 5% CO_2_. Cells were then fixed, permeabilized, and stained using eBioscience BrdU Flow kit according to manufacturer’s directions.

### RT-qPCR

RNA was isolated from marrow cells using RNeasy Mini Kit (Qiagen) or TRIzol (Invitrogen). cDNA was prepared using SuperScriptII (Invitrogen) and amplified using PowerSybr Green Master (Applied Biosystems) and HT7900 sequence detection system (Applied Biosystems) per manufacture specifications. Primers used were as follows: mIL-1β (for-AAGGCTGCTTCCAAACCTTTGACC; rev-ATACTGCCTGCCTGAAGCTCTTGT); mIL-6 (for-TACCACTTCACAAGTCGGAGGCTT; rev-CAATCAGAATTGCCATTGCACAAC); mIL-10 (for-GCAGGACTTTAAGGGTTACTTGGG; rev-CCTTGATTTCTGGGCCATGCTTCT); mIL-12p40 (for-ATGTGGGAGCTGGAGAAAGACGTT; rev-ATCTTCTTCAGGCGTGTCACAGGT); mTNFα (for-TCTCAGCCTCTTCTCATTCCTGCT; rev-GCCATTTGGGAACTTCTCATCCCT); mIfhβ (for-TGAACTCCACCAGCAGACAGTGTT; rev-TCAAGTGGAGAGCAGTTGAGGACA); mIfnγ (for-TGCCAAGTTTGAGGTCAACAACCC; rev-TTTCCGCTTCCTGAGGCTGGATT); mIL-2Rα (for-GCAAGAGAGGTTTCCGAAGA; rev-CGATTTGTCATGGgAGTTGC); mCXCL10 (for-GCTGCAACTGCATCCATATC; rev-GTGGCAATGATCTCAACACG); mCSF2 (for-AGGGTCTACGGGGCAATTT; rev-ACAGTCCGTTTCCGGAGTT); mFasL (for-TTTAACAGGGAACCCCCACT; rev-GATCACAAGGCCACCTTTCT); mIfna (for-GACTTTGGATTCCCGCAGGAGAAG; rev-CTGCATCAGACAGCCTTGCAGGTC); mIfnab (for-CCTGCTGGCTGTGAGGAAAT; rev-CTCACTCAGACTTGCCAGCA); mHmgb1 (for-GTTCTGAGTACCGCCCCAAA; rev-GTAGGCAGCAATATcCTTCTC); mIL-4 (for-TTGAACGAGGTCACAGGAGA; rev-AAATATGCGAAGCACCTTGG); mIL-17a (for-CGCAAAAGTGAGCTCCAGA; rev-TGAGCTTCCCAGATCACAGA); mIL-1rn (for-GTGAGACGTTGGAAGGCAGT; rev-GCATCTTGCAGGGTCTTTTC); mTNFSF11 (for-GACTCCATGAAAACGCAGGT; rev-CCCACAATGTGTTGCAGTTC); mIL-13 (for-TGTGTCTCTCCCTCTGACCC; rev-CACACTCCATACCATGCTGC); mIL-1a (for-AGCGCTCAAGGAGAAGACC; rev-CCAGAAGAAAATGAGGTCGG); mIL-27 (for-GTGACAGGAGACCTTGGCTG; rev-AGCTCTTGAAGGCTCAGGG); mCSF1 (for-CCAGGATGAGGACAGACAGG; rev-GGTAGTGGTGGATGTTCCCA); mCCL2 (for-AGGTCCCTGTCATGCTTCTG; rev-GGGATCATCTTGCTGGTGAA); mIL-23a (for-TTGTGACCCACAAGGACTCA; rev-AGGCTCCCCTTTGAAGATGT); mIfna13 (for-CTTTGGATTCCCACAGGAGA; rev-TTCCATGCAGCAGATGAGTC); mIfna2 (for-GCAGATCCAGAAGGCTCAAG; rev-GGTGGAGGTCATTGCAGAAT); mGAPDH (for-TGCAGTGGCAAAGTGGAGAT; rev-TTTGCCGTGAGGAGTCATA); hIFNγR1 (for-CCAGGGTTGGACAAAAAGAA; rev-CGGGATCATAATCGACTTCC); hIFNγR2 (for-TGACAATGCCTTGGTTTCAA; rev-ATCAGCGATGTCAAAGGGAG); mCamp (for-GCTGTGGCGGTCACTATCAC; rev-TGTCTAGGGACTGCTGGTTGA); mTmsb10 (for-CCGGACATGGGGGAAATCG; rev-CCTGTTCAATGGTCTCTTTGGTC); mTmsb4x (for-ATGTCTGACAAACCCGATATGGC; rev-CCAGCTTGCTTCTCTTGTTCA); mAnxa6 (for-CACAGGGTGCCATGTACCG; rev-TCCTGATTTGCGTCAAACTCTG); mAnxa1 (for-ATGTATCCTCGGATGTTGCTGC; rev-TGAGCATTGGTCCTCTTGGTA); mAnxa5 (for-ATCCTGAACCTGTTGACATCCC; rev-AGTCGTGAGGGCTTCATCATA); mLamc1 (for-TGCCGGAGTTTGTTAATGCC; rev-CTGGTTGTTGTAGTCGGTCAG); mLamb1 (for-GAAAGGAAGACCCGAAGAAAAGA; rev-CCATAGGGCTAGGACACCAAA); hCasp1 (for-GCTTTCTGCTCTTCCACACC; rev-CACATCACAGGAACAGGCAT); hHMGB1 (for-TCGGGAGGAGCATAAGAAGA; rev-CTCTTTCATAACGGGCCTTG); hHPRT (for-TGACCTTGATTTATTTTGCATACC; rev-CGAGCAAGACGTTCAGTCCT); hGAPDH (for-TGATTCCACCCATGGCAAATTCC; rev-GCTCCTGGAAGATGGTGATGGATT). GAPDH or hHPRT served as reference control and differences in mRNA expression levels were calculated as fold-changes by the ΔΔCt method.

### Enzyme-linked immunosorbent assay

We plated TIRAP- or vector-transduced marrow cells at a concentration of 2 x 10^6^ cells/ml in serum free DMEM medium supplemented with 1% BSA and cultured them overnight. We recovered the supernatants and centrifuged (3500rpm, 5 min) and assayed for Il10 and Hmgb1. Serum from peripheral blood collected at endpoint from mice transplanted with TIRAP- or vector-transduced HSPC was used to measure Ifnγ. Mouse IL-10 ELISA Ready-Set-Go!, Mouse IFNγ ‘Femto-HS’ High Sensitivity ELISA and HMGB1 ELISA kits from eBioscience were used according to manufacturer’s instructions.

### Homing Assay

3×10^5^ retrovirally transduced marrow cells were transplanted into lethally irradiated recipient mice. Mice were euthanized 22 hours post transplant and GFP content was analyzed by flow cytometry. Total number of transduced GFP+ marrow cells was determined. Homing efficiency of progenitor cells was determined by comparing the proportion of GFP+ colonies formed in colony forming assay before transplantation and 22 hrs after transplantation. GFP+ colonies were counted using a Axiovert S100 fluorescent microscope (Zeiss, Oberkochen, Germany). Homing efficiency was calculated using the following formula:

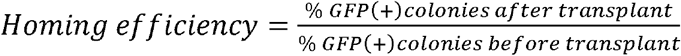

### Endothelial cells cytotoxicity assay

Human Umbilical Vein Endothelial cells (HUVECs) were grown in MCDB131 medium supplemented with 20% FBS, 50μg/ml Heparin and 50μg/ml endothelial cell growth supplement. 8 x 10^4^ HUVECs were seeded in 200μl in 96 well plates, 24 hours before starting treatment with Hmgb1. To measure the cytotoxic effect of HMGB1, we treated HUVECs with 200ng/ml HMGB1 or 200ng/ml HMGB1+ 500μg/ml Glycyrrhizin or PBS (as control) in minimal medium (MCDB 131 supplemented with insulin, sodium selenite, transferrin and 0.1% BSA). After 24 hours of treatment, the cell viability was measured by adding 44μM of Resazurin solution to the wells and incubated for 4 hours. After incubation, the fluorescence was measured at 560nm excitation. The % Viability was calculated as:

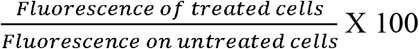

### RNA-seq

Total RNA samples were checked using Agilent Bioanalyzer RNA nanochip or Caliper GX HT RNA LabChip. Samples that passed quality check were arrayed into a 96-well plate. Following this, polyA+ RNA was purified using the 96-well MultiMACS mRNA isolation kit on the MultiMACS 96 separator (Miltenyi Biotec, Germany) from total RNA with on column DNaseI-treatment as per the manufacturer’s instructions. The eluted polyA+ RNA was ethanol precipitated and resuspended in 10μL of DEPC treated water with 1:20 SuperaseIN (Life Technologies, USA). Double-stranded cDNA was synthesized from the purified polyA+ RNA using the Maxima H Minus First Strand cDNA Synthesis Kit (Thermo Fisher Scientific Inc., USA) and random hexamer primers. Quality passed cDNA plate was fragmented by Covaris LE220 for 2×65 seconds at “Duty cycle” of 30%. The paired-end sequencing library was prepared following the BCCA Genome Sciences Center paired-end library preparation plate-based library construction protocol on a Biomek FX robot (Beckman-Coulter, USA). Briefly, the cDNA was subject to end-repair, and phosphorylation by T4 DNA polymerase, Klenow DNA Polymerase, and T4 polynucleotide kinase respectively in a single reaction, followed by cleanup using magnetic beads and 3’ A-tailing by Klenow fragment (3’ to 5’ exo minus). After cleanup, adapter ligation was performed. The adapter-ligated products were purified using magnetic beads, then UNG digested and PCR-amplified with Phusion DNA Polymerase (Thermo Fisher Scientific Inc., USA) using Illumina’s PE primer set in a single reaction, with cycle condition 37°C 15min, 98°C 1min followed by 13 cycles of 98°C 15 sec, 65°C 30 sec and 72°C 30 sec, and then 72°C 5min. The PCR products were purified and size selected using magnetic beads, checked with Caliper LabChip GX for DNA samples using the High Sensitivity Assay (PerkinElmer, Inc. USA) and quantified with the Quant-iT dsDNA HS Assay Kit using Qubit fluorometer (Invitrogen). Libraries were normalized and pooled. The final concentration was double checked and determined by Qubit dsDNA HS Assay for Illumina Sequencing.

### Gene expression and pathway Analysis

RNA-Seq data was aligned using GSNAP (version 2013-10-28) (47), using the mm10_jg-e71 reference and with command-line arguments: ‘_novelsplicing 1 –max-mismatches 10 –use-splicing’. We used RNA-SeQC (48) to gather quality metrics for the RNA-Seq libraries. Expression quantification was performed with sailfish (v0.9.0) (49), using RefSeq gene models downloaded as GTF from the UCSC genome browser on 2014-08-21, with gene models from non-standard chromosome sequences removed. Both isoform- and gene-specific quantifications were generated, and both raw estimated counts, as well as transcripts-per-million (TPM) estimates were used in downstream analysis. The differential expression analysis was performed using DESeq2 (v1.10.1) (50), using an FDR-adjusted p-value cut-off of 0.1 to identify differentially expressed genes between the two groups. Pathway analysis was performed using the differentially expressed gene dataset obtained from RNA-Seq as input in the Ingenuity Pathways Analysis (IPA) software, a web-delivered application that enables biologists to discover, visualize and explore therapeutically relevant networks significant to their experimental results, such as gene expression array data sets (https://www.qiagenbioinformatics.com/products/ingenuity-pathway-analysis/). The identified genes were mapped to genetic networks available in the Ingenuity database and then ranked by score and assigned a *P* value. The score was defined by the probability that a collection of genes equal to or greater than the number in the respective network could be achieved by chance alone.

### GSEA

Gene set enrichment analysis software (Broad Institute) _ENREF_48 was used to analyze data from previously published microarray data sets. The Supplementary Table 4 lists the genes included in the IFNγ and IL10 gene sets.

### Statistics

Statistical analyses were performed using Prism 6.01 (GraphPad Software Inc., La Jolla, CA). A two-sided unpaired Students’t-test was employed. ANOVA analysis was used for multiple group comparisons. The error bars represent SEM in all figures. For survival analysis, the Mantel-Cox test was used. For all data, statistical significance was considered at *P ≤* 0.05.

### Study approval

All animal protocols were approved by the Animal Care Committee of the University of British Columbia (Vancouver, British Columbia).

## Author Contributions

AK conceived the study. AG, RI and AK designed experiments; AG, RI, MF, PU, LC, JW, JLam, and ML performed experiments; AG, RI, JLi, JP and AK analyzed data; AG, RI and AK wrote the manuscript.

## Acknowledgments

This work was supported by grants to AK from a Terry Fox New Frontiers Program Project and the Canadian Institutes of Health Research, as well as funding from the BC Cancer Foundation through the Leukemia and Myeloma Program of BC. AK is the recipient of the John Auston BC Cancer Foundation Award.

