## Supplemental Figures (1-7) for "TIRAP drives myelosuppression through an Ifnγ-Hmgb1 axis that disrupts the marrow microenvironment"

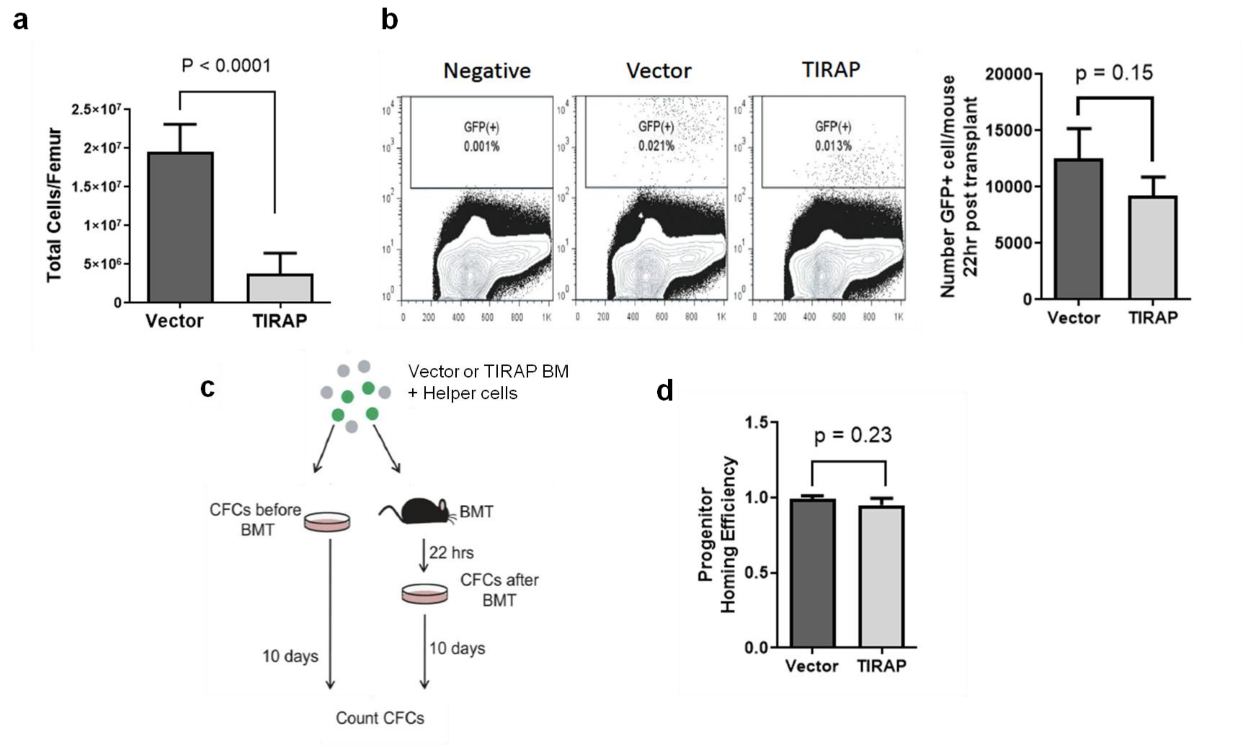


**Supplemental Fig. 1: Marrow failure is not due to a homing defect**

(a) Trypan blue counts of total cells (live + dead) in the femurs of mice transplanted with TIRAP (n = 15) or Vector (n = 6) transduced HSPCs at endpoint. (b) Flow cytometric analysis of BM collected from mice transplanted with Vector (n = 4) or TIRAP (n = 4) transduced marrow 22 hours post-transplant and quantification of the total number of transduced cells that homed to the marrow per mouse. (c) Schematic illustrating progenitor homing assay. (d) Quantification of progenitor homing efficiency. Homing efficiency was calculated using the following formula: $Homing\ efficiency = \frac{\%\ GFP(+) colonies\ after\ transplant}{\%\ GFP(+) colonies\ before\ transplant}$.

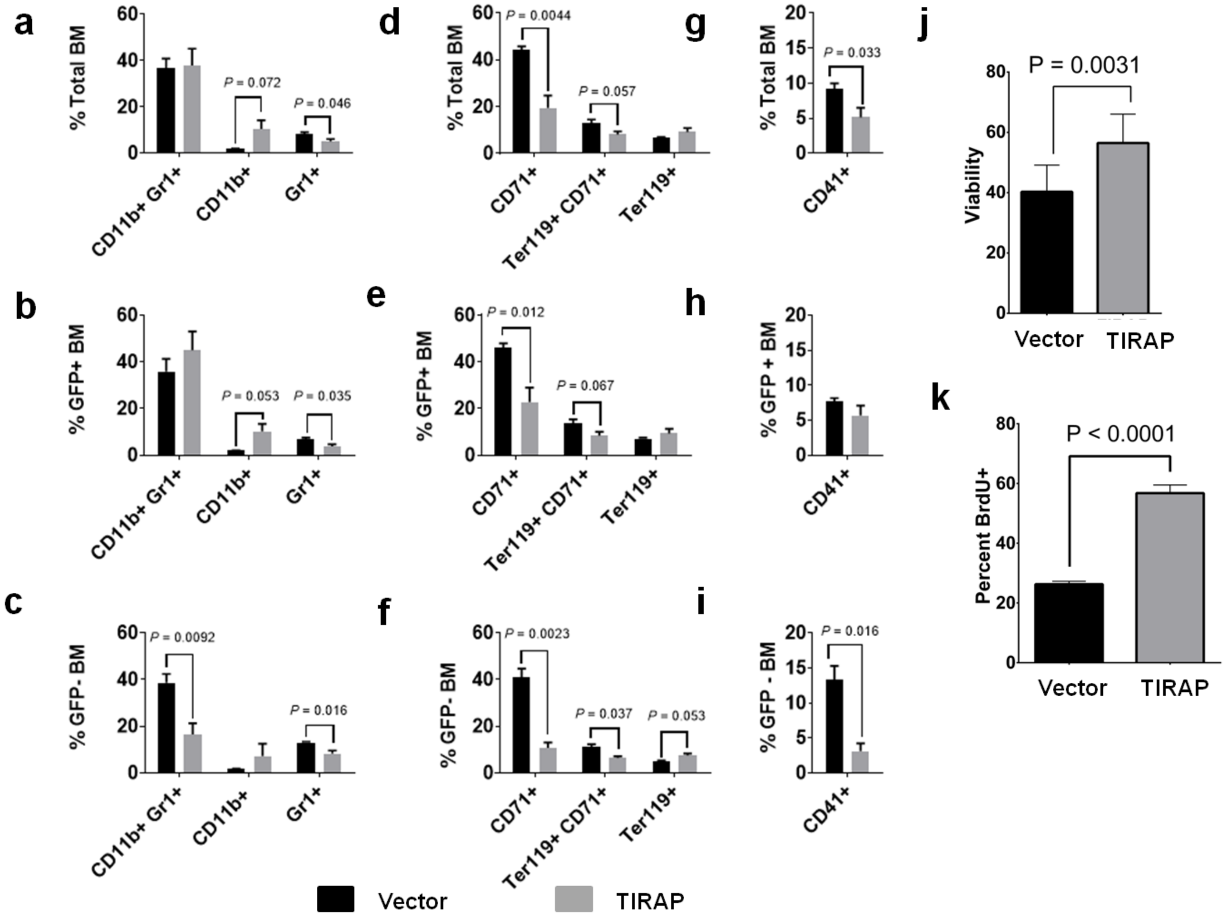


**Supplemental Fig. 2: Constitutive TIRAP expression disrupts normal hematopoiesis in a cell nonautonomous manner.**

Immunophenotyping of myeloid (a-c) erythroid (d-f) and megakaryocytic (g-i) lineages from the marrow of wild-type mice transplanted with Vector- (n = 3) or TIRAP- (n = 6) transduced wild-type HSPC at 3-4 weeks post transplant. Total HSPC (upper), as well as the contribution of transduced (GFP+; middle) and competitor cells (GFP- ; lower) are presented. Cell viability was assessed by Annexin V/PI staining (j), and cell cycling was measured by BrdU incorporation (k) in TIRAP- or Vector-transduced HSPC *in vitro*.

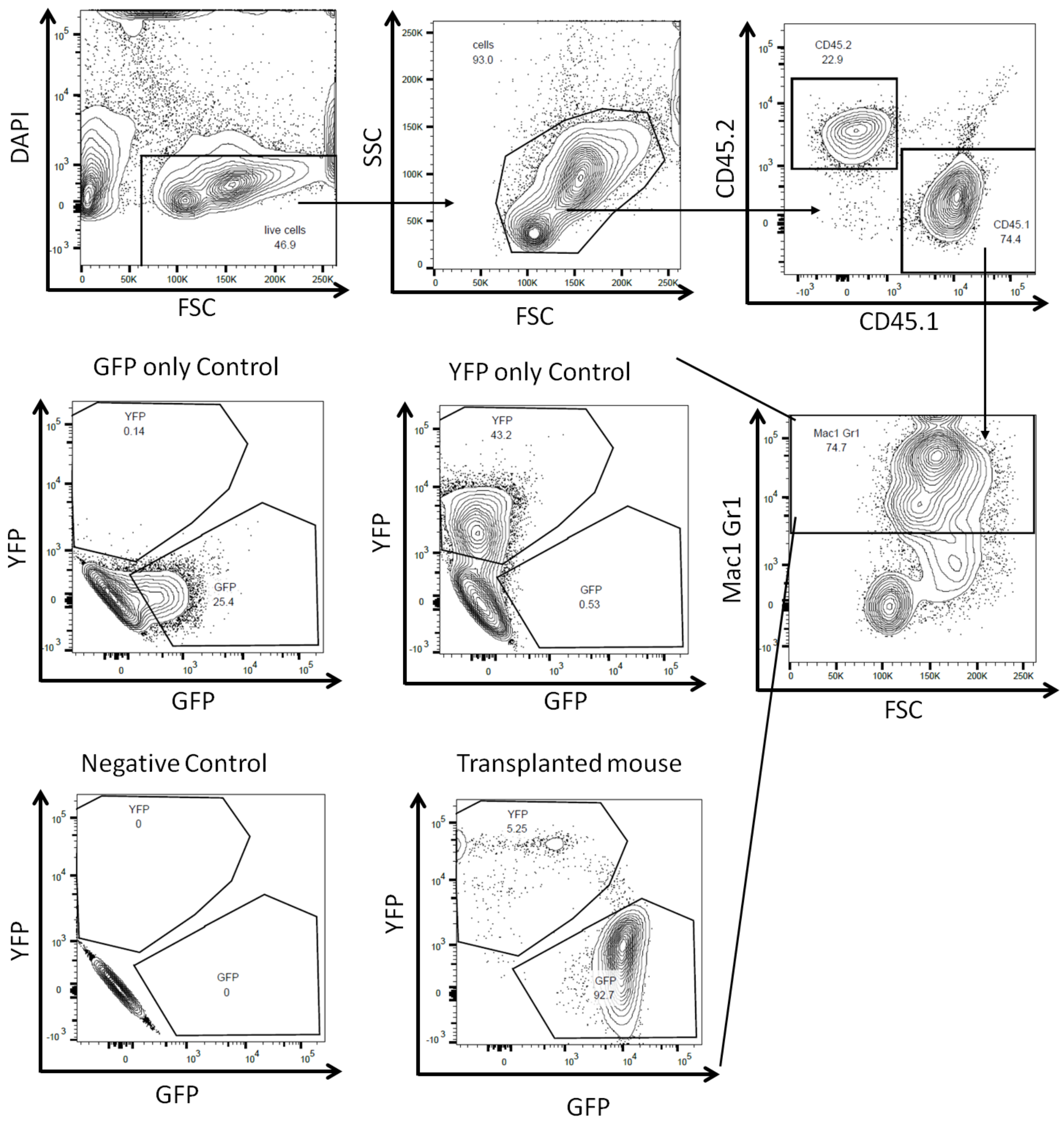


**Supplemental Fig. 3: HSPC gating strategy for determining marrow contribution of GFP- and YFP-labeled cells in chimeric mice**

Gating strategy for analysis of contributions of GFP- and YFP-labeled HSPC to hematopoietic reconstitution in the myeloid cell populations. The CD45.1 gate was used to identify donor cells in the marrow of the transplanted mouse. $Mac1^{+}Gr1^{+}$ cells were identified as myeloid cell populations. The GFP and YFP positive cells were gated using negative, GFP- and YFP- only controls.

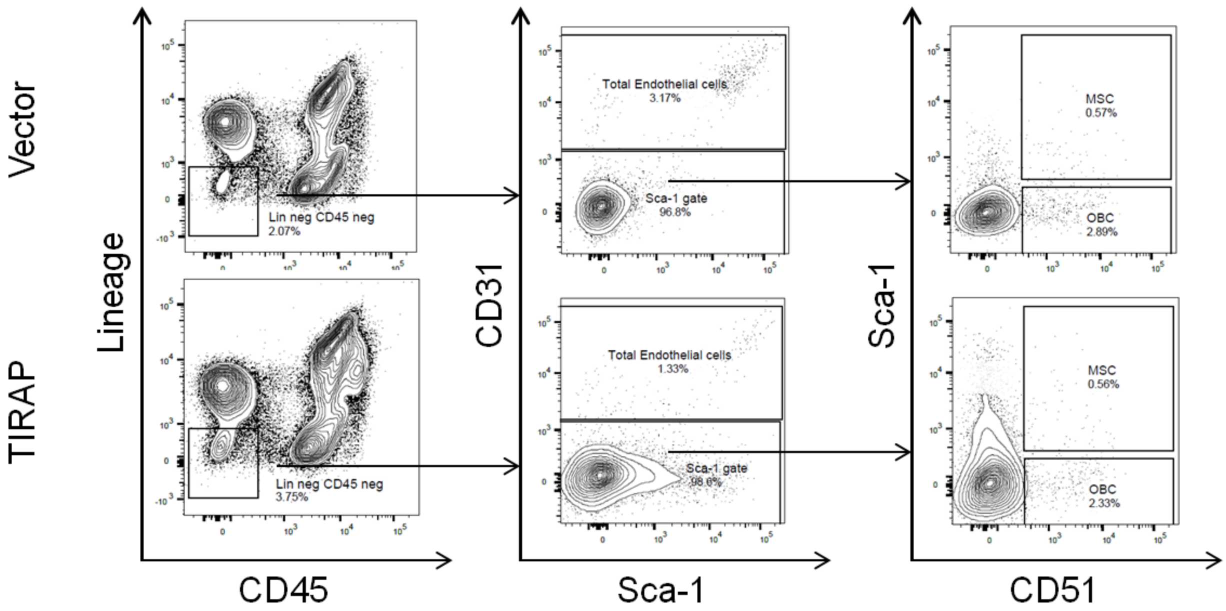


**Supplemental Fig 4: Bone marrow stromal cell gating strategy**

Staining of marrow stromal cells was performed as previously described (32). Total endothelial cells were identified as lin$^-$CD45$^-$CD31$^+$, mesenchymal stromal cells were identified as lin$^-$CD45$^-$ Sca-1$^+$CD51$^+$, and osteoblastic cells were identified as lin$^-$CD45$^-$Sca-1$^-$CD51$^+$.

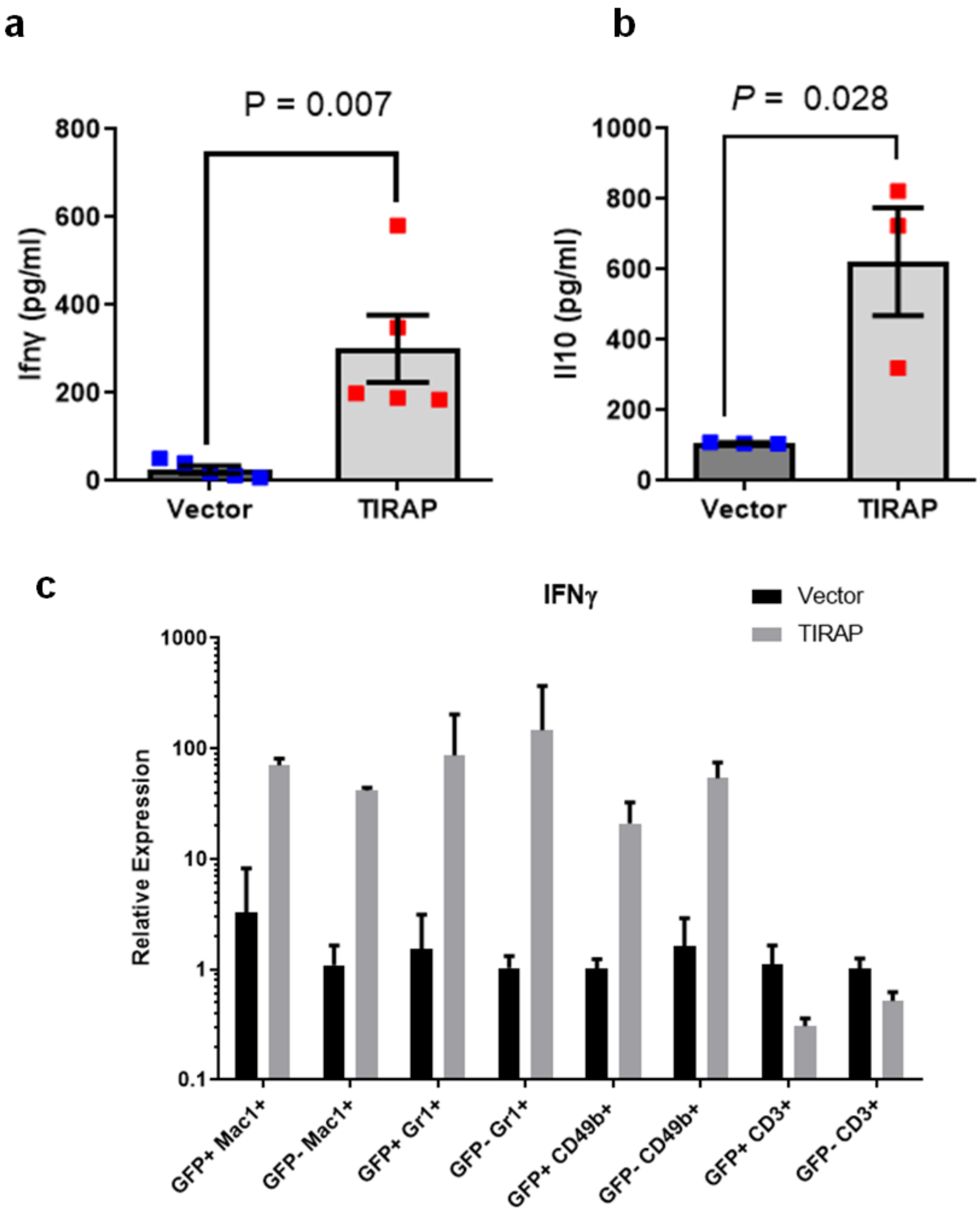


**Supplemental Fig. 5: Constitutive expression of TIRAP induces Ifnγ and Il-10**

(a) Concentration of Ifnγ in peripheral blood serum from mice transplanted with TIRAP- or Vector-transduced HSPC at experimental end-point. (b) Concentration of Il10 detected in conditioned medium from HSPC transduced with TIRAP or Vector after 24 hours of culture. (c) Ifnγ levels as measured by intracellular qPCR at experimental endpoint in myeloid cells ($Mac1^+$ and $Gr1^+$) NK cells ($CD49b^+$) and T cells ($CD3^+$) in the bone marrow of wild-type mice transplanted with Vector- or TIRAP- transduced wild-type HSPC. Ifnγ levels were measured in transduced (GFP+) and competitor cells (GFP-).

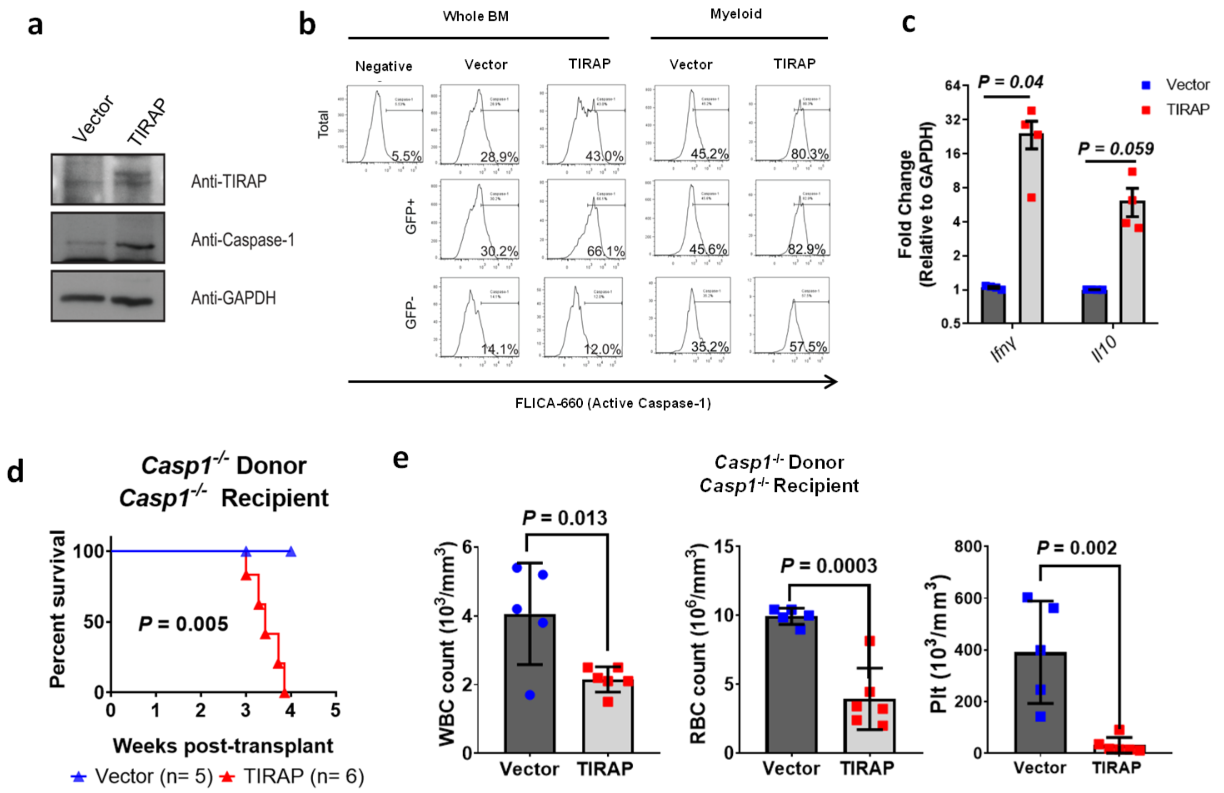


**Supplemental Fig. 6: Caspase-1 is not required for TIRAP-induced BMF**

(a) Immunoblots for Caspase-1 from wild-type HSPC transduced with TIRAP or Vector. (b) Intracellular flow cytometric analysis of activated Caspase-1. CD11b$^+$ cells were selected to measure the Active Caspase-1 within myeloid populations. Marrow from 4 mice was pooled and analyzed for activated Caspase-1. (c) RT-qPCR measuring *Ifnγ* and *Il10* in *Casp1*$^{-/-}$ HSPC transduced with TIRAP or Vector. (d) Kaplan-Meier survival curves for *Casp1*$^{-/-}$ mice transplanted with *Casp1*$^{-/-}$ HSPC transduced with TIRAP (n = 6) or Vector (n = 5). (e) Complete blood counts from *Casp1*$^{-/-}$ mice transplanted with *Casp1*$^{-/-}$ HSPC transduced with TIRAP or Vector.

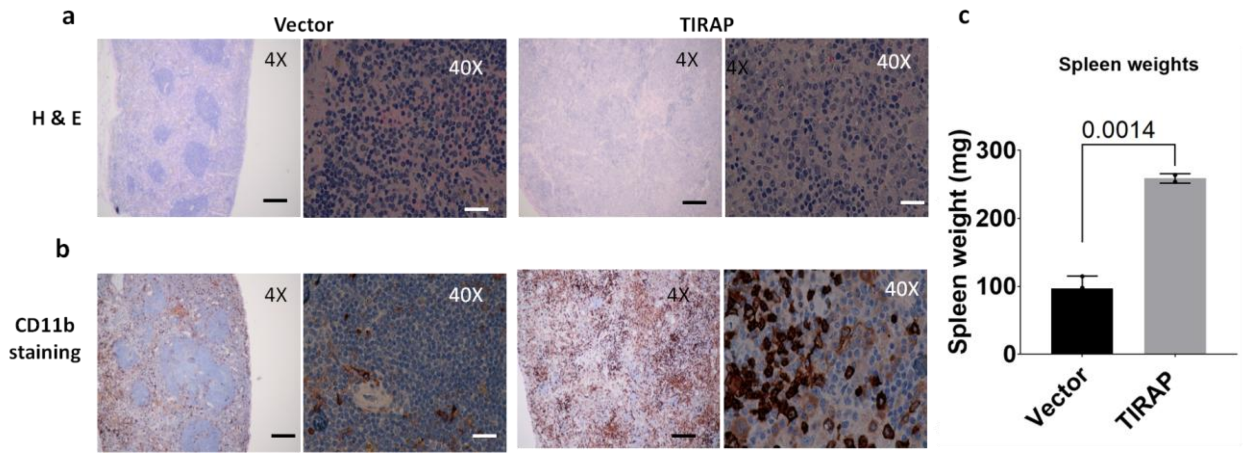


**Supplemental Fig. 7: Loss of Ifnγr leads to splenic infiltration of myeloid cells upon constitutive expression of TIRAP.**

(a) Hematoxylin and eosin-stained sections of the spleen of Ifnγr mice transplanted with TIRAP-transduced HSPC lacking Ifnγr show disruption of the splenic structures compared to vector controls. (b) CD11b-stained spleen sections show increase infiltration of myeloid cells in Ifnγr mice transplanted with TIRAP-transduced HSPC lacking Ifnγr compared to Vector. Scale bar: 200 µm for 4x magnification and 20 µm for 40X magnification. (c) Spleen weight at experimental endpoint of Ifnγr mice transplanted with Vector- and TIRAP- transduced HSPC lacking Ifnγr.
