## Supplemental table 1 for "TIRAP drives myelosuppression through an Ifnγ-Hmgb1 axis that disrupts the marrow microenvironment"

Table S1. Top 20 cytokines predicted to regulate differential gene expression

| Upstream Regulator | Predicted Activation State | p-value of overlap |
|---|---|---|
| IFNG | Activated | 4.29E-22 |
| IFNB1 | Activated | 1.17E-15 |
| IFNL1 | Activated | 4.24E-14 |
| IL10 | NA | 2.48E-13 |
| IL6 | Activated | 4.73E-13 |
| TNF | Activated | 6.10E-13 |
| IL4 | NA | 4.01E-12 |
| IL1B | Activated | 5.45E-12 |
| IL17A | Activated | 3.08E-11 |
| IL1RN | Inhibited | 5.66E-10 |
| TNFSF11 | Activated | 9.74E-10 |
| IFNA1/IFNA13 | Activated | 9.98E-10 |
| IL13 | NA | 2.80E-09 |
| IL1A | Activated | 3.91E-09 |
| IFNA2 | Activated | 5.88E-08 |
| IL27 | Activated | 3.49E-07 |
| CSF1 | NA | 4.33E-07 |
| CCL2 | NA | 4.44E-07 |
| IL23A | NA | 4.55E-07 |
| IL18 | Activated | 6.17E-07 |

NA- not assessable
