## Supplemental table 3 for "TIRAP drives myelosuppression through an Ifnγ-Hmgb1 axis that disrupts the marrow microenvironment"

**Table S3: Pathways Significantly activated downstream of TIRAP**

| Ingenuity Canonical Pathways | -log(p-value) |
|---|---|
| Communication between Innate and Adaptive Immune Cells | 9.43E00 |
| Role of Pattern Recognition Receptors in Recognition of Bacteria and Viruses | 7.74E00 |
| Altered T Cell and B Cell Signaling in Rheumatoid Arthritis | 6.47E00 |
| TREM1 Signaling | 4.25E00 |
| phagosome formation | 4.25E00 |
| Dendritic Cell Maturation | 4.21E00 |
| Differential Regulation of Cytokine Production in Macrophages and T Helper Cells by IL-17A and IL-17F | 4.19E00 |
| Granulocyte Adhesion and Diapedesis | 4.15E00 |
| Differential Regulation of Cytokine Production in Intestinal Epithelial Cells by IL-17A and IL-17F | 3.87E00 |
| NF-κB Signaling | 3.87E00 |
| T Helper Cell Differentiation | 3.85E00 |
| Crosstalk between Dendritic Cells and Natural Killer Cells | 3.52E00 |
| Agranulocyte Adhesion and Diapedesis | 3.47E00 |
| Systemic Lupus Erythematosus Signaling | 3.4E00 |
| Role of Macrophages, Fibroblasts and Endothelial Cells in Rheumatoid Arthritis | 3.34E00 |
| Pathogenesis of Multiple Sclerosis | 3.16E00 |
| Role of IL-17A in Psoriasis | 3.16E00 |
| Role of Hypercytokinemia/hyperchemokinemia in the Pathogenesis of Influenza | 3.07E00 |
| Complement System | 2.66E00 |
| LXR/RXR Activation | 2.46E00 |
| Chemokine Signaling | 2.38E00 |
| B Cell Development | 2.25E00 |
| Xenobiotic Metabolism Signaling | 2.24E00 |
| Allograft Rejection Signaling | 2.18E00 |
| Toll-like Receptor Signaling | 2.15E00 |
| PCP pathway | 2.08E00 |
| IL-12 Signaling and Production in Macrophages | 2.01E00 |
| Dermatan Sulfate Biosynthesis (Late Stages) | 1.92E00 |
| Hepatic Fibrosis / Hepatic Stellate Cell Activation | 1.87E00 |
| Role of Cytokines in Mediating Communication between Immune Cells | 1.87E00 |
| Autoimmune Thyroid Disease Signaling | 1.87E00 |
| Atherosclerosis Signaling | 1.87E00 |
| Graft-versus-Host Disease Signaling | 1.72E00 |
| Methylglyoxal Degradation III | 1.68E00 |
| Activation of IRF by Cytosolic Pattern Recognition Receptors | 1.63E00 |
| **HMGB1 Signaling** | 1.63E00 |
| Bladder Cancer Signaling | 1.5E00 |
| Melatonin Degradation III | 1.47E00 |
| Role of IL-17F in Allergic Inflammatory Airway Diseases | 1.46E00 |
| FcγRIIB Signaling in B Lymphocytes | 1.42E00 |
| Dermatan Sulfate Biosynthesis | 1.39E00 |
| Actin Cytoskeleton Signaling | 1.36E00 |
| Regulation of the Epithelial-Mesenchymal Transition Pathway | 1.36E00 |
| PI3K Signaling in B Lymphocytes | 1.33E00 |
| GM-CSF Signaling | 1.32E00 |
