## Supplemental table 4 for "TIRAP drives myelosuppression through an Ifnγ-Hmgb1 axis that disrupts the marrow microenvironment"

**Table S4: IFNγ & IL-10 gene sets**

**IFNγ stimulated genes**

| | | | | | | |
|---|---|---|---|---|---|---|
| ABCD3 | CD38 | FGFR1 | IFIT3 | ISG15 | SECTM1 | TP63 |
| ATF3 | CEACAM1 | GBP1 | IFIT5 | ISG20 | STAT1 | TYMP |
| BAX | CFLAR | GBP2 | IFITM1 | ITGB7 | STX11 | UBE4B |
| BRCA2 | CXCL10 | IFI16 | IFITM3 | KNTC1 | TAP1 | VCAM1 |
| CASP1 | CXCL11 | IFI27 | IFNG | LAMP3 | TMOD1 | WARS |
| CASP10 | CXCL9 | IFI30 | IGF1R | MX1 | TNF | |
| CASP8 | DNAJA2 | IFI35 | IGFBP4 | NAMPT | TNFAIP2 | |
| CCL5 | EIF2AK2 | IFI44 | IL10RA | OPTN | TNFAIP6 | |
| CCL8 | FAS | IFI6 | IL12RB1 | PML | TNFSF10 | |
| CCNA1 | FASLG | IFIT2 | IRF1 | RAB7L1 | TOP1 | |

**IL-10 signalling pathway genes**

| | | | | | |
|---|---|---|---|---|---|
| BLVRA | IL10 | IL1A | STAT1 | STAT4 | STAT6 |
| BLVRB | IL10RA | IL6 | STAT2 | STAT5A | TNF |
| HMOX1 | IL10RB | JAK1 | STAT3 | STAT5B | |
